# Histone H2A ubiquitination resulting from Brap loss of function connects multiple aging hallmarks and accelerates neurodegeneration

**DOI:** 10.1101/2020.10.22.341784

**Authors:** Y. Guo, A. A. Chomiak, Y. Hong, C. C. Lowe, W-C. Chan, J. Andrade, H. Pan, X. Zhou, E. Berezovski, E. S. Monuki, Y. Feng

**Author notes:** Kite Pharma, Santa Monica, CA 90404, U.S.A.

## Abstract

Aging is an intricate process that is characterized by multiple hallmarks including stem cell exhaustion, genome instability, epigenome alteration, impaired proteostasis, and cellular senescence. While each of these traits is detrimental at the cellular level, it remains unclear how they are interconnected to cause systemic organ deterioration. Here we show that abrogating Brap, a BRCA1 associated protein important for neurogenesis, results in cellular senescence with persistent DNA double-strand breaks and elevation of histone H2A mono- and poly-ubiquitination (H2Aub). The high H2Aub initiates histone proteolysis, leading to both epigenetic alteration and proteasome overflow. These defects induce neuroinflammation, impair proteostasis, accelerate neurodegeneration, and substantially shorten lifespan in mice carrying Brap deletions in the brain. We further show H2Aub is also increased in human brain tissues of Alzheimer’s disease. These data together suggest that chromatin aberrations mediated by H2Aub act as a nexus of multiple aging hallmarks and promote tissue-wide degeneration.

## INTRODUCTION

Aging is a natural process that connects birth and death. The time-dependent decline in organ function during aging is the major risk factor for many diseases including neurodegenerative disorders (NDs). Yet, the etiology of aging is complex and multifactorial. It is associated with multiple cellular and molecular hallmarks, such as stem cell exhaustion, genome instability, epigenetic alteration, deregulated nutrient sensing, mitochondrial dysfunction, cellular senescence, and loss of proteostasis (Lopez-Otin et al., 2013). For neurodegeneration, additional hallmark defects in neuronal calcium homeostasis and neuronetwork activity, as well as increased glial cell activation and neuroinflammation are also essential contributors to the physical and functional deterioration of the brain (Mattson and Arumugam, 2018). Aging at the tissue, organ, and system levels does not simply result from the dysfunction of any individual cellular or molecular process. To cause progressive and irreversible tissue-or organ-wide degeneration, multiple aging promoting factors must act in concert. However, the mechanism underlying the interconnections among various aging hallmarks is poorly understood, forming a major barrier to combating aging-associated diseases.

Among various aging hallmarks, genome instability poses a lifelong threat to all living cells and can result in increased somatic mutations that have been shown to accumulate with age in most organs (Lombard et al., 2005; Soares et al., 2014; White and Vijg, 2016). Besides a clear link to carcinogenesis, loss of genome stability has also been implicated in neurodegeneration (Bushman et al., 2015; Madabhushi et al., 2014; Mitra et al., 2019; Rulten and Caldecott, 2013; Shanbhag et al., 2019; Thadathil et al., 2019), and there has been long-established evidence for significant levels of DNA damage even in non-dividing cells, such as neurons in the brain (Lodato et al., 2015; Lu et al., 2004; Rutten et al., 2007; Suberbielle et al., 2013). Recent whole genome sequencing analyses have further demonstrated that somatic mutations increase with age in both bulk tissues and single neurons of the human brain (Hoang et al., 2016; Lodato et al., 2018). Moreover, mutations of genes important for genome maintenance and DNA repair cause progeroid syndromes and often manifest with both premature aging and neurodegeneration (Choy and Watters, 2018; Coppede and Migliore, 2010; Kakigi et al., 1992; Weidenheim et al., 2009). Thus, genotoxicity is a universal insult that not only is concomitant with old age but also drives aging-associated diseases. Nevertheless, the mechanism by which neuronal DNA damage non-autonomously affects many cells across the brain tissue to promote neurodegeneration remains elusive.

A plausible mechanism for DNA damage to induce tissue-wide degeneration is via cellular senescence, a stress response that occurs when cells experience telomere erosion, chronic oncogenic stimulation, persistent DNA damage, oxidative and metabolic stresses, or mitochondrial dysfunction (Hayflick, 1965; Hernandez-Segura et al., 2018; McHugh and Gil, 2018; Munoz-Espin and Serrano, 2014; Rodier and Campisi, 2011). Although originally defined by a permanent replication arrest, cellular senescence is also characterized by a set of distinctive phenotypes, including activation of senescence-associated β-galactosidase (SA-β-gal), expression of a cyclin-dependent kinase inhibitor (CKI) p16^Ink4a^, alterations of global epigenetic and metabolic profiles, resistance to apoptosis, increases in autophagy, and implementation of a complex and multicomponent secretome known as the senescence-associated secretory phenotypes (SASP) (Campisi and Robert, 2014; Coppe et al., 2008; Munoz-Espin and Serrano, 2014; Rodier and Campisi, 2011). Through SASP, senescent cells produce a myriad of pro-inflammatory cytokines, chemokines, growth factors, and proteases (Kuilman and Peeper, 2009; Lasry and Ben-Neriah, 2015). These secreted molecules act through autocrine and paracrine signaling to not only reinforce the senescence state, but also non-autonomously affect neighboring cells in the tissue. Remarkably, senescence-like phenotypes have also been observed in neurons with tau aggregation and neurofibrillary tangles of the Alzheimer’s disease (AD) brains (Musi et al., 2018). Therefore, it is possible that neurons, which are replication-incompetent by nature, can acquire the additional characteristics of senescent cells upon chronic DNA damage to promote neuroinflammation and neurodegeneration.

We identified BRAP as a key partner of NDE1 that is essential for human brain development and brain genome protection (Alkuraya et al., 2011; Bakircioglu et al., 2011; Houlihan and Feng, 2014; Lanctot et al., 2013). BRAP/Brap is a ubiquitously expressed cytoplasmic protein originally cloned through its interaction with the BRCA1 tumor suppressor (Li et al., 1998). It is also a Ras responsive ubiquitin E3 ligase that modulates the sensitivity of MAPK signaling (Matheny et al., 2004). Our previous studies showed that this role of Brap in neural progenitor cells (NPCs) was context-dependent and that Brap E3 ligase could act on other E3 ligases with wide nuclear substrates (Lanctot et al., 2017; Lanctot et al., 2013). These distinctive features allow Brap to serve as a liaison between the cell’s microenvironment and cell nucleus to coordinate cell signaling, differentiation, and genome stability. Moreover, the essential role of Brap goes beyond brain development. In a recent genome-wide association study of 500,000 genotyped individuals along with information on their parents’ age and cause of death, the *BRAP* gene locus was linked to the human lifespan (Timmers et al., 2019), suggesting an interplay of BRAP function and human aging.

By analyzing Brap null (Brap^-/-^) and cerebral cortical neural progenitor cell (NPC) Brap conditional knockout (Brap^cKONPC^) mice, we have shown that Brap loss of function (LOF) selectively affected the neuronal fate restrictive cell cycle of multipotent NPCs, exhibiting accelerated G1 phase, abrogated G1/S checkpoint, and prolonged S and G2 phases. The skewed cell cycle kinetics impeded the neurogenesis of NPCs and resulted in reduced generation of cerebral cortical neurons (Lanctot et al., 2017). We now demonstrate that besides attenuating neurogenesis, Brap LOF also results in cellular senescence, affecting both NPCs and neocortical neurons. We show that senescence in Brap^-/-^ and Brap^cKONPC^ cells is accompanied by persistent DNA double-strand breaks (DSBs) and elevations of Brca1 as well as histone H2A ubiquitination (H2Aub), catalyzed by Brca1 E3 ubiquitin ligase. Notably, the high H2Aub initiates histone proteolysis, causing not only global epigenetic alteration but also increased proteasome burden. In the cortical tissue of Brap^cKONPC^ mice, the backlog of polyubiquitinated proteins and SASP together result in neuroinflammation, neurodegeneration, and midlife mortality. We also demonstrate that H2Aub, the hallmark phenotype of Brap LOF, is elevated in cortical tissue of patients with Alzheimer’s disease (AD). These results suggest that histone H2Aub is a key node connecting multiple aging hallmarks, highlighting an essential impact of chromatin aberrations on NDs.

## RESULTS

### Brap LOF results in accelerated cellular senescence with persistent DSBs

To understand the mechanism by which Brap LOF skews cell cycle kinetics and impedes the differentiation of NPCs, we isolated NPCs from Brap^-/-^ embryos (before embryonic lethality by E13.5) and studied them in culture as neurospheres. Although Brap^-/-^ NPCs grew at a rate comparable to their wild-type (WT) counterparts in the first 4-5 days *in vitro* (div), they abruptly ceased proliferation after 7-8 days (Figure 1A). This growth arrest was not accompanied by increased apoptosis (Figure 1B), which suggested that the mutant NPCs reached their replicative lifespan due to cellular senescence.

**FIGURE 1.**
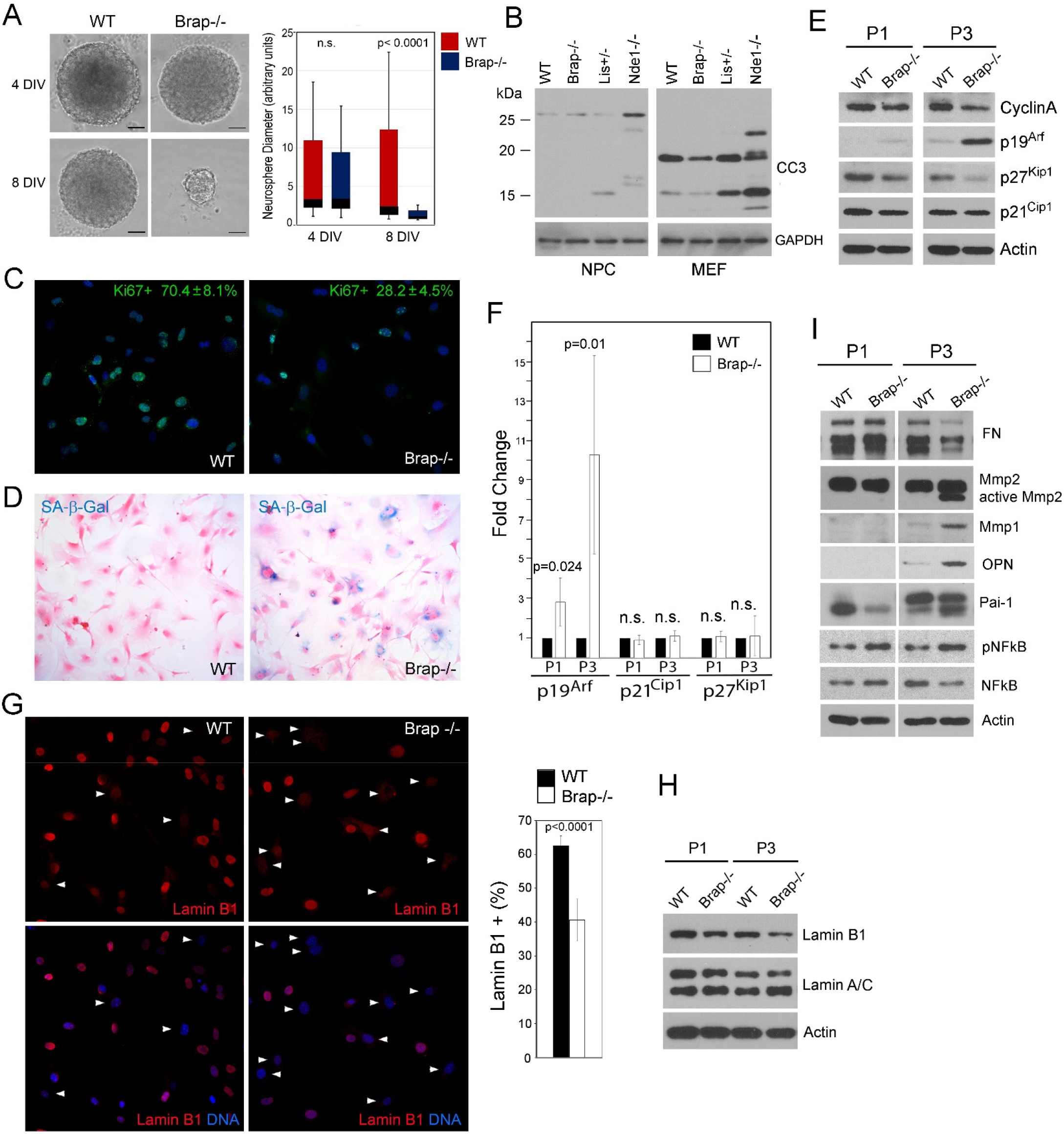
BRAP LOSS OF FUNCTION (LOF) RESULTS IN ACCELERATED CELLULAR SENESCENCE. (A) Primary Brap^-/-^ neural progenitor cells (NPC) cease growing in culture after 5-7 days in vitro (DIV). Shown are representative images of WT and Brap^-/-^ neurospheres as well as box and whisker plots (Median ± SD) of their diameter distribution at 4 DIV and 8 DIV, respectively. (B) Cleaved caspase 3 (CC3) immunoblotting of total protein extracts of primary NPCs and MEFs, respectively. Lis1^+/-^ and Nde1^-/-^ cells were used for CC3+ control as these cells were known to have high levels of apoptosis. (C) Representative Ki67 immunofluorescence images of WT and Brap^-/-^ MEFs at P2. The % of Ki67+ cells (Mean ± SD) is indicated. (D) Senescence-associated β−gal assay of MEFs at P2, which indicates β-galactosidase activity (blue) in higher fraction of Brap^-/-^ cells. Cells were also counter-stained with eosin to view cell density and cell shape. Note the enlarged cell body of β−gal+ cells. (E, F) Immunoblotting and quantification (Mean ± SD) of G1/S CKIs in total protein extracts from MEFs at P1 (Passage 1) and P3, respectively. (G) Representative Lamin B1 immunofluorescence images and percentage (Mean ± SD) of Lamin B1+ MEFs at P2. Cells showing significant reduction of nuclear Lamin B1 are indicated by arrowheads. (H) Immunoblotting of total protein extracts, showing reduced Lamin B1 in Brap^-/-^ relative to WT MEFs at both P1 and P3. (I) Immunoblotting of total protein extracts from MEFs at P1 and P3, showing increased SASP molecules in Brap^-/-^ MEFs at P3. Note the increased Mmps and decreased Mmp substrate fibronectin (FN) in Brap^-/-^ MEFs. Nuclear DNA was stained with Hoechst 33342. Bars: 100 um or as indicated.

To determine cellular senescence we turned to mouse embryonic fibroblast (MEFs) as they are better established for senescence analyses. We isolated MEFs from Brap^-/-^ and WT embryos and cultured them following the 3T3 cell protocol (Todaro and Green, 1963). As expected, a greater fraction of Brap^-/-^ MEFs showed multiple essential characteristics of cellular senescence as early as passage 2 (P2). These include an enlarged cell body, high senescence-associated beta-galactosidase (SA-β–gal) activity, loss of the Ki67 proliferation marker, and upregulation of p19^Arf^, a CKI encoded by the p16^Ink4a^ locus that is alternatively transcribed and can act together with p16^Ink4a^ to induce cellular senescence (Capparelli et al., 2012; Haber, 1997; Quelle et al., 1995) (Figure 1C-F). We further validated the senescence phenotype of Brap LOF by examining the loss of Lamin B1, another characteristic of cellular senescence (Freund et al., 2010). Both immunofluorescence (IF) and immunoblotting (IB) analyses demonstrated that Lamin B1 was downregulated much more rapidly in Brap^-/-^ than WT MEFs from P1 to P3 (Figure 1G, H). To acquire the ultimate proof of cellular senescence, we assessed several known SASP molecules, including Mmp1, Mmp2, Serpin1 (Pai1), and Osteopontin/Spp1 (OPN) (Aoshiba et al., 2013; Castello et al., 2017; Coppe et al., 2010; Flanagan et al., 2018; Ghosh and Capell, 2016; Pazolli et al., 2012; Rao and Jackson, 2016; Vaughan et al., 2017). We found that all of them were elevated and the Mmp substrate fibronectin reduced in Brap^-/-^ MEFs along with increased phosphorylation of NFkB p65^S536^, a chief mediator for SASP (Salminen et al., 2012) (Figure 1I). The multiple essential features of cellular senescence shown by Brap^-/-^ cells demonstrates unambiguously that accelerated cellular senescence is a consequence of Brap LOF. These data also suggest that the attenuated neurogenesis of Brap^-/-^ NPCs in cortical development was caused by a pro-senescence fate change.

While senescence is essentially a stress response to a variety of insults, we found that cellular senescence caused by Brap LOF was associated with persistent DNA damage responses (DDRs). First, we detected significant elevation of phospho-p53, -Atm, -Atr, and 53BP1 in Brap^-/-^ MEFs as early as in P1 (Figure 2A-C). In addition, a large fraction of Brap^-/-^ MEFs showed persistent DNA damage foci with high levels of γH2A.X, the hallmark of DNA double-strand breaks (DSBs), and 53BP1, a marker for DSB repair (Panier and Boulton, 2014; Thiriet and Hayes, 2005)(Figure 2D-F). Notably, the large γH2A.X and 53BP1 foci were in both proliferating (Ki67+) cells and cells that had ceased dividing (Ki67-) (Figure 2D and E, high magnification panels), indicating that DSBs persisted in Brap^-/-^ cells that had undergone replication arrest. Furthermore, we found that the rapid progression to senescence of Brap^-/-^ MEFs coincided with increased levels of γH2A.X ubiquitination (Figure 2F). While mono-ubiquitination of histone H2A is a DNA damage response, the ubiquitinated moiety of γH2A.X has been shown to mark non-apoptotic DSBs induced by oxidative DNA damage (Luczak and Zhitkovich, 2018). Consistent with this, we found that ubiquitin-modified γH2A.X (γH2A.Xub) in Brap^-/-^ cells was further enhanced by H_2_O_2_ (Figure 2F), suggesting that γH2A.Xub is an apoptosis-evading and senescence-prone DNA damage response of Brap LOF.

**FIGURE 2.**
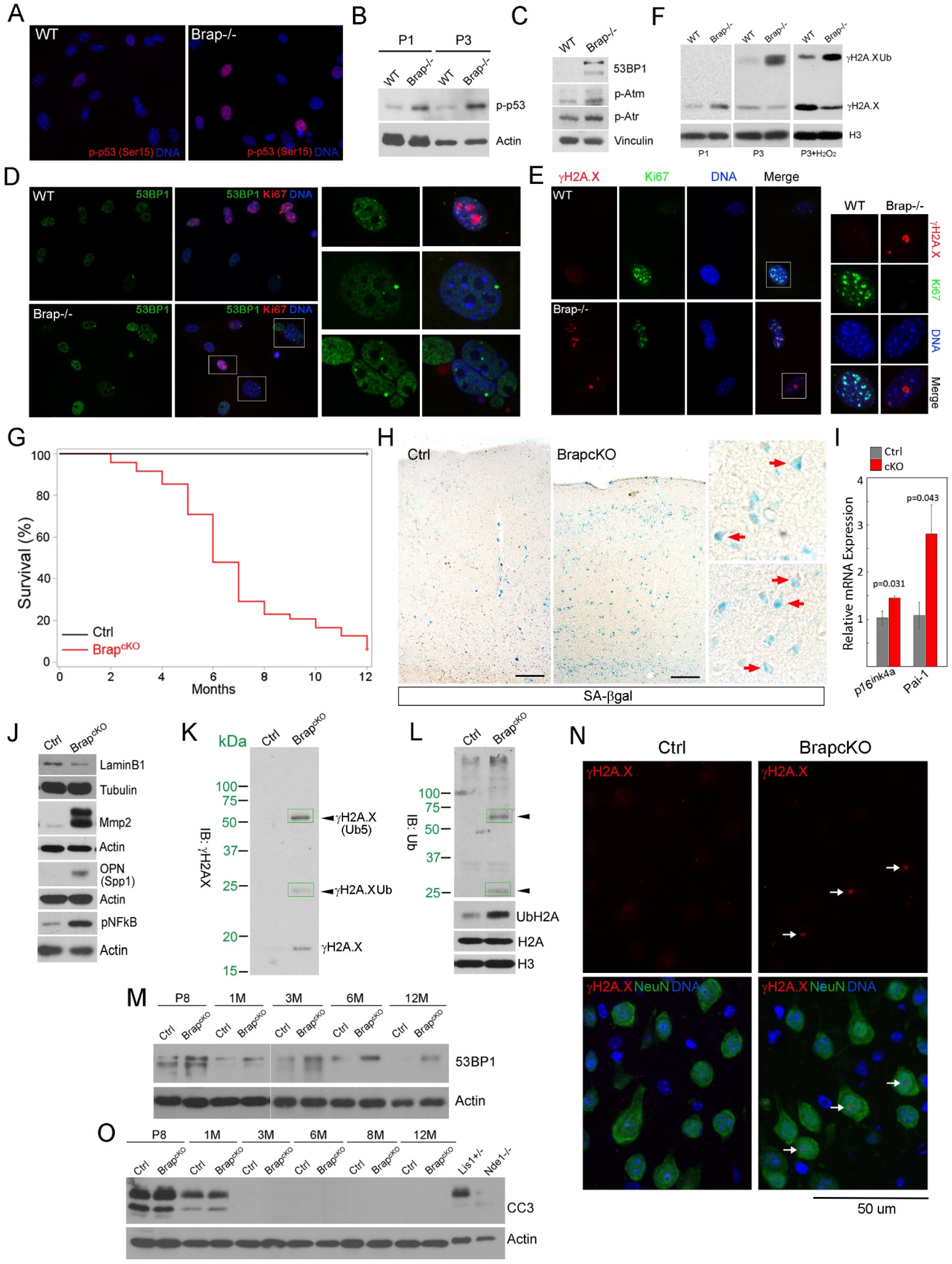
INCREASED DSB AND DDR ARE ASSOCIATED WITH CELLULAR SENESCENCE IN BRAP DEFICIENT CELLS AND CEREBRAL CORTICES. (A) Representative images of phospho-p53 immunofluorescence staining of WT and Brap^-/-^ MEFs at P2. (B) Immunoblotting of total protein extracts from WT and Brap^-/-^ MEFs at P1 and P3, respectively. (C) Immunoblotting of DNA damage response (DDR) proteins in WT and Brap^-/-^ MEFs at P1. (D) Representative images of double immunofluorescence staining of WT and Brap^-/-^ MEFs at P3 with antibodies against Ki67 (red) and 53BP1 (green). (E) Representative images of double immunofluorescence staining of WT and Brap^-/-^ MEFs at P3 with antibodies against Ki67 (green) and γH2A.X (red). (F) Immunoblotting analyses of histone extracts of MEFs at P1 or P3, showing increased γH2A.X and γH2A.X mono-ubiquitination in Brap^-/-^ MEFs. (G) Kaplan-Meier curve shows significantly shortened lifespan of Brap^cKONPC^ mice (n=46) relative to their littermate control mice (n=29). (H) Representative images of senescence-associated β−gal analysis of cerebral cortical sections from Brap^cKONPC^ and control mice at 3 months of age. Note the pyramidal neuronal morphology of some SA-βgal+ cells (red arrows) in high magnification views. (I) RT-qPCR shows increased expression of p16^Ink4a^ and Pai-1 (plasminogen activator inhibitor-1/Serpine1) in 3-month old Brap^cKONPC^ cortices (Mean ± SD; n=4 biological replicates). (J) Immunoblotting of cerebral cortical total protein extracts, demonstrating higher levels of senescence and SASP-like changes in the cortical tissue of Brap^cKONPC^ relative to littermate control mice. (K, L) Immunoblotting of histone extracts from cortical tissue of 3-month old Brap^cKONPC^ and control mice, showing elevated γH2A.X as well as the presence of mono- and poly-γH2A.X ubiquitination in Brap^cKONPC^ mice (boxed bands). K and L are the same immunoblot that was first probed by anti-γH2A.X, stripped, and re-probed by anti-ubiquitin. (M) Immunoblotting of total protein extracts from cortical tissue of Brap^cKONPC^ and control mice, showing persistent elevation of 53BP1 in Brap^cKONPC^ cortices from P8 (postnatal day 8) to 12M (12 months). N. Representative images of γH2A.X-NeuN double immunohistological staining of cerebral cortical sections from 4-month old WT or Brap^cKONPC^ mice. The presence of γH2A.X immunoreactivity in Brap^cKONPC^ neurons is indicated by arrows. (O) Immunoblotting of total protein extracts from cortical tissue of Brap^cKONPC^ or control mice at various ages, showing that Brap LOF does not increase cleaved caspase 3, an apoptosis marker. Embryonic Lis1^+/-^ and Nde1^-/-^ cortical tissues were used as positive controls for the presence of apoptosis. Nuclear DNA was stained with Hoechst 33342. Bars: 100 um or as indicated.

### Cellular senescence and DSBs in cerebral cortical tissue underlie shortened lifespan of Brap^cKONPC^ mice

We subsequently found that the accelerated cellular senescence and DSBs can also occur in the brain, causing shortened lifespan of mice in which Brap was conditionally ablated in NPCs of the of the dorsal telencephalon by the Emx1-Cre. In these Brap conditional knockout (cKO) mice (Brap^flox Emx1Cre+^, referred to as Brap^cKONPC^ hereafter), Cre-mediated deletion results in Brap LOF in glutamatergic neurons, astroglia, and oligodendrocytes of the cerebral cortex (Gorski et al., 2002). We followed a cohort of adult Brap^cKONPC^ mice (n= 46) and their Brap^flox/WT^ or Brap^flox/flox^ Cre-control littermates (n=29) over 12 months. We found that most Brap^cKONPC^ mice die between 4 and 8 months with a median lifespan of 6 months (Figure 2G), whereas the average lifespan of mice is 2 years. The mutant mice lost weight, became lethargic, and showed slow or labored breathing when death became imminent (Supplementary Video). Therefore, Brap LOF in the cerebral cortex can accelerate aging and shorten the lifespan.

With midlife mortality, Brap^cKONPC^ mice showed high levels of cellular senescence in the cerebral cortical tissue. We first examined SA-β-gal activity in cortical sections. We found it was not only higher in Brap^cKONPC^ than in control mice but also shown by many mutant cells with pyramidal neuron morphology (Figure 2H, arrows). This suggested that, despite being replication-incompetent, neurons can undergo additional senescence-associated changes due to Brap LOF. Further corroborating the increased cellular senescence, the cortical tissue of Brap^cKONPC^ mice showed upregulated p16^Ink4^, decreased Lamin B1, and elevation of SASP factors including Serpin1 (Pai-1), Mmp2, Ssp1 (OPN), and phospho-NFkB (Figure 2I and J).

Similar to the observations in Brap^-/-^ MEFs, increased cell senescence in Brap^cKONPC^ cortical tissue was associated with the presence of DSBs. We found that γH2A.X was not only elevated in the cortical tissue of Brap^cKONPC^ compared to control mice but also mono- and poly-ubiquitinated, showing a dominant pool of approximately 60 kDa that is equivalent to the molecular weight of γH2A.X with Penta-ubiquitin conjugations (Figure 2K, L). Such dual PTM in response to DSBs with simultaneous phosphorylation and poly-ubiquitination of H2A.X has not been reported before but appears to be characteristic of the DDR of Brap^cKONPC^ cortical tissue. Coinciding with increased γH2A.X, Brap^cKONPC^ cortical tissues also showed higher levels of 53BP1 (Figure 2M). Our immunohistological (IH) analysis showed that γH2A.X immunoreactivities were restricted to the nucleus of a substantial fraction of cortical neurons (NeuN+) in the brain of Brap^cKONPC^ mice but was neither in mutant glia (NeuN-) nor in the brain of WT mice (Figure 2N). Given that we did not observe tumors or elevated apoptosis in the brain Brap^cKONPC^ mice (Figure 2O), cellular senescence is likely the main consequence of persistent DSBs and accounts for the premature mortality of Brap^cKONPC^ mice.

### Transcriptomic profiles of Brap^cKONPC^ cortices reveal SASP candidates, immune activation, and impaired synaptic signaling

Compared to apoptosis, senescence following unsuccessful DSB repair is more deleterious *in vivo*, as senescent cells are stably viable and can chronically influence neighboring cells by secreting soluble molecules or by altering cell surface and extracellular matrix (ECM) proteins. Although many senescence associated secretory molecules have been identified in various culture and *in vivo* models, the SASP profile of cellular senescence in the brain has not been investigated. We thus sought to assess senescence associated brain secretome by identifying molecules with increased expression from Brap LOF in cortical tissue. RNA sequencing (RNA-seq) was carried out with cerebral cortical total transcripts of Brap^cKONPC^ and littermate control mice of 3 months old. We identified 811 differentially expressed genes (DEGs) between the Brap^cKONPC^ and the control group, and of these, 373 genes were significantly upregulated by Brap LOF (Figure 3A, Supplementary Table 1). Consistent with the pro-inflammatory feature of cellular senescence, we found 67 of the upregulated genes encode molecules associated with cells of the immune system and/or regulators of the innate immunity (Figure 3B). The upregulation of these genes indicates inflammatory responses in the mutant brain tissue. Moreover, 80 upregulated genes encode secreted molecules, which represent the cerebral cortical secretome of Brap^cKONPC^ mice. Increased expression of these molecules can pose non-cell autonomous effects via SASP. These SASP candidate genes include not only immune active molecules and proteases but also diverse neuropeptide neurotransmitters and peptide hormones with potent activities in neural networking and cerebrovascular regulation (Figure 3C). An additional 83 upregulated genes encode plasma membrane and ECM proteins that interface cells with their environment (Figure 3D). Increased expression of these genes can alter intercellular communication and tissue homeostasis. The most significantly upregulated genes in Brapc^KONPC^ cortices also include regulators for neurotransmitter metabolism, fatty acid and cholesterol metabolism, protein synthesis and sorting, as well as calcium homeostasis (Figure 3A, Supplementary Table 1). These molecular activities can have broad impacts, resulting in a global alteration in brain immunity and neural function.

**FIGURE 3.**
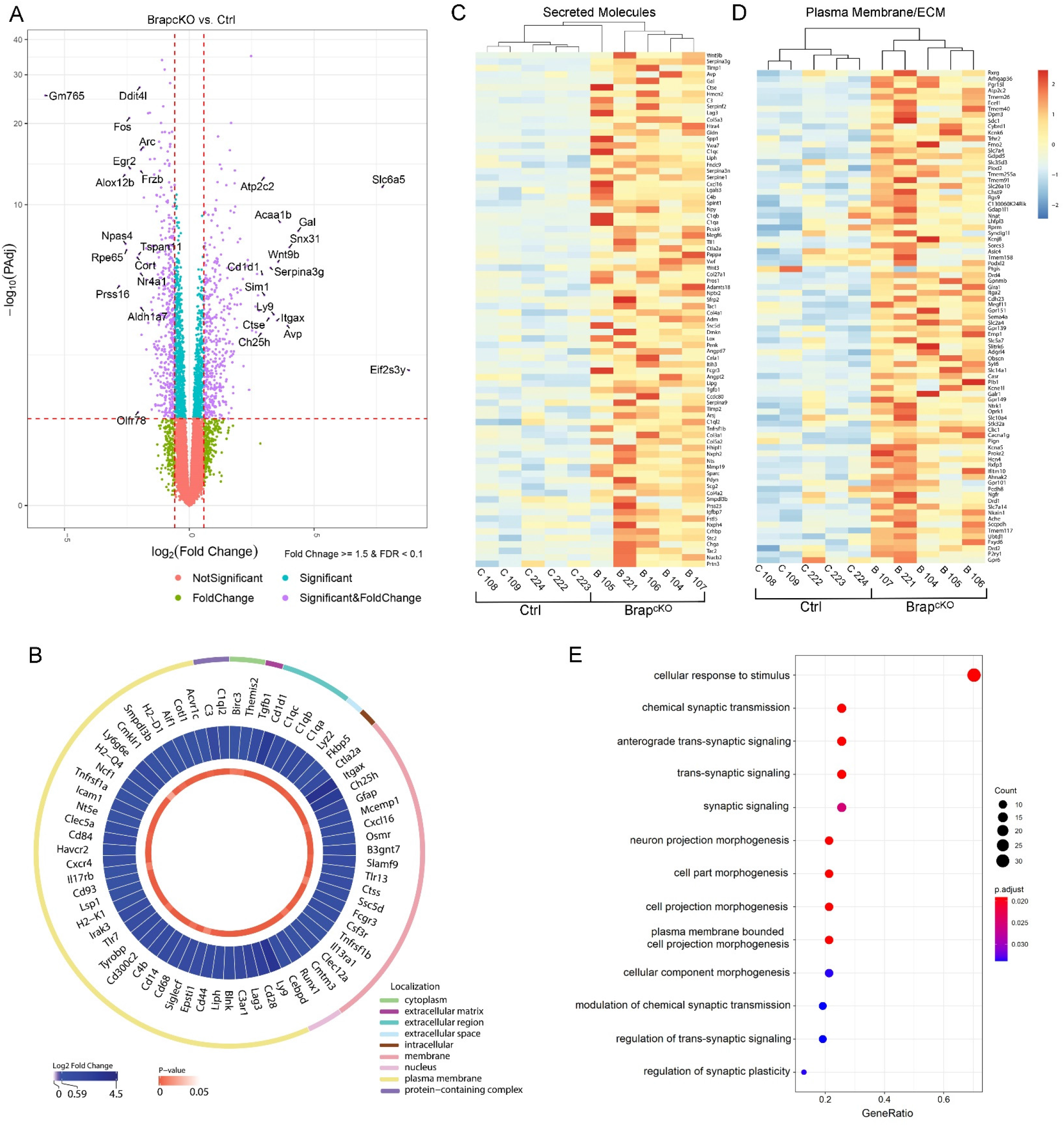
DIFFERENTIAL GENE EXPRESSION CAUSED BY BRAP LOF REVEALS INCREASED IMMUNE ACTIVITIES, SASP PROFILES, AND COMPROMISED SYNAPTIC FUNCTION IN CEREBRAL CORTICAL TISSUE. (A) Volcano plot shows differentially expressed genes (DEGs) between Brap^cKONPC^ and control cortical tissues. (B) Circos plot presentation of upregulated genes associated with immune cells and innate immunity regulations in Brap^cKONPC^ relative to control cerebral cortices. (C) Heatmap shows increased expression of genes encoding secreted molecules in Brap^cKONPC^ relative to control cerebral cortices. (D) Heatmap shows increased expression of genes encoding plasma membrane and extracellular matrix molecules in Brap^cKONPC^ relative to control cerebral cortices. (E) Gene ontology (GO) enrichment analysis of the top 50 down-regulated genes (by adjusted p-value) in Brap^cKONPC^ relative to control cortical tissues. Shown are dot plots of biological processes affected by Brap LOF. Note that synaptic signaling, transmission, and cellular response are significantly compromised in Brap^cKONPC^ cortices.

Brap LOF also resulted in decreased expression of 438 genes in the cortical tissue (Supplementary Table 1). Notably, the most significant downregulation occurred in genes that play key roles in regulating energy homeostasis, neuronal activity, synaptic plasticity, neuronal excitatory-inhibitory balance, Wnt signaling, retinoid metabolism, as well as the production, secretion, and activity control of neuropeptides or neurotransmitters (Figure 3A). We performed Gene Ontology (GO) enrichment analysis of the 50 most significantly downregulated genes in Brap^cKONPC^ cortices. The GO terms identified in the BP category include impaired biological functions mainly in synaptic transmission as well as in cellular responses to stimuli and neuronal morphogenesis (Figure 3E), suggesting declining neuronal activity and plasticity. Overall, our transcriptomic data provide additional support for an impact of neuronal DSBs and cellular senescence on neuroinflammation and neural dysfunction of Brap^cKONPC^ mice.

### Neuroinflammation and neurodegeneration in Brap^cKO^ brains

Inflammatory reactions in the cortical tissue are not only characteristics of cellular senescence but also contributors to neurodegeneration. We thus asked whether neurodegeneration underlies the premature mortality of Brap^cKONPC^ mice. As expected, neuroinflammatory and neurodegenerative phenotypes were readily detectable in Brap^cKONPC^ cortices starting at 3 months and progressing rapidly with age. First, hyperphosphorylation of the microtubule-associated protein tau became significantly increased in the cortical tissue of Brap^cKONPC^ mice at 3 months and was further elevated as the mutant mice grew older (Figure 4A and B). Increased tau phosphorylation in Brap^cKONPC^ cortices was found on multiple residues including T181, T217, S396, and S416 (Figure 4A). The hyperphosphorylation on these residues is known to be associated with insoluble tau aggregates and neurodegenerative diseases. As the presence of these human tauopathy-like changes in Brap^cKONPC^ mice was at ages when mortality started to occur, it supported the notion that the mutant mice died of neurodegenerative brain dysfunction.

**FIGURE 4.**
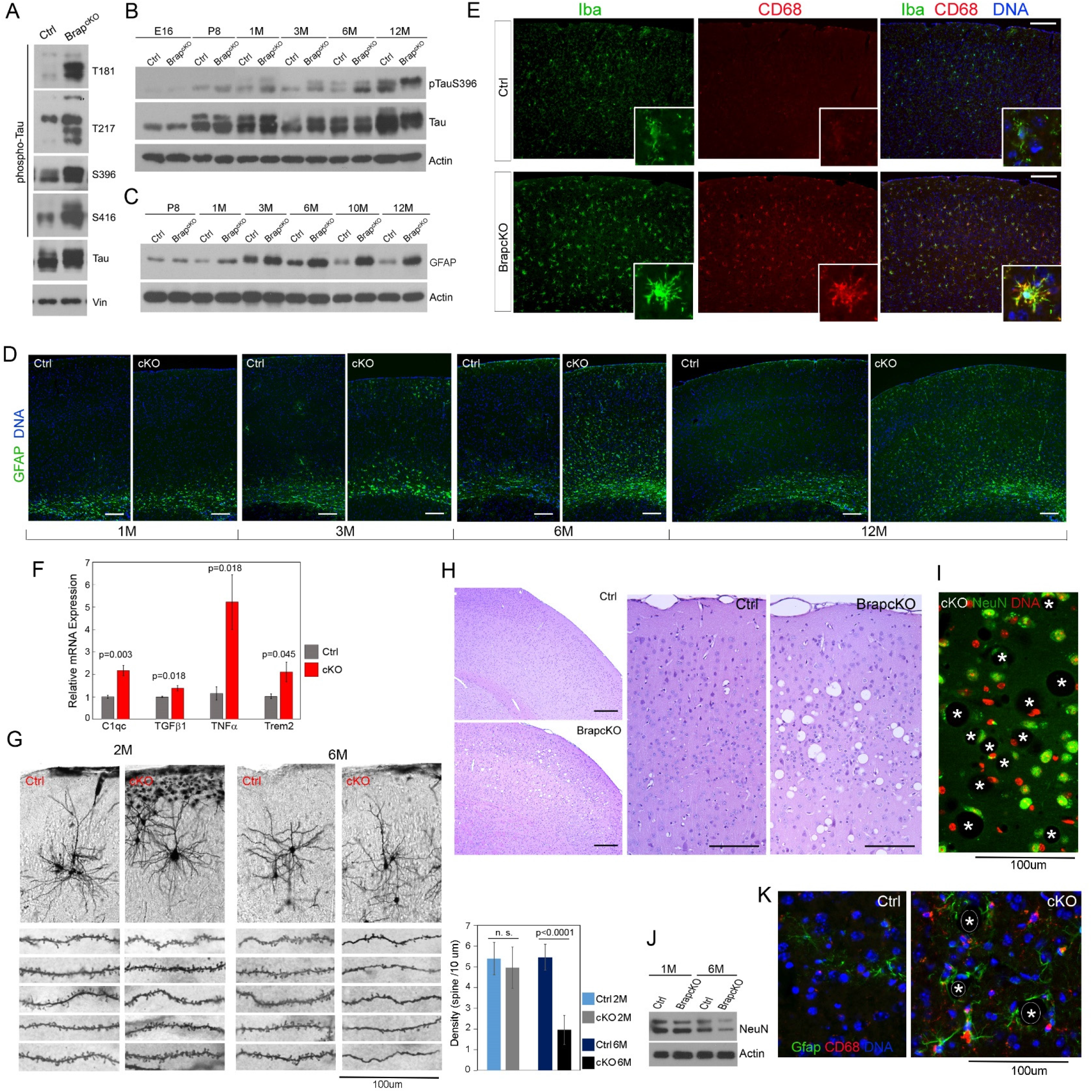
BRAP LOF IN CEREBRAL CORTICAL NPCS RESULTS IN NEUROINFLAMMATION AND ACCELERATED NEURODEGENERATION. (A) Immunoblotting of cerebral cortical total protein extracts from 3-month old Brap^cKONPC^ and control mice with various anti-phospho-tau antibodies. (B) Immunoblotting of cortical total proteins extracts shows age-dependent progression of tau hyper-phosphorylation (on S396) in Brap^cKONPC^ cortices. (C, D) Gfap immunohistological and immunoblotting analyses show age-dependent progression of astrogliosis in Brap^cKONPC^ cortices. (E) Double immunohistological staining with anti-Iba (green) and anti-CD68 (red) antibodies shows marked increase in microglia activation in Brap^cKONPC^ cortices at 3 months. Representative images are shown. F. RT-qPCR analyses of selected neuroinflammatory genes in 3-month cortical tissues (Mean ± SE; n = 4-8 biological replicates). (G) Representative images and quantification (Mean ± SD) of Golgi-cox analyses, showing significantly reduced density of dendritic spines in cortical layer 2/3 pyramidal neurons of Brap^cKONPC^ mice at 6 months. H. H&E stained brain sections of 10 months old Brap^cKONPC^ and control mice, showing spongiform encephalopathy in Brap^cKONPC^ cortical gray matter. (I) A representative NeuN immunohistological image shows reduced neuronal density and dystrophic or distorted neurons adjacent to spongiosis (positions of vacuoles are marked by asterisks). (J) Immunoblotting of cortical total protein extracts, demonstrating decreased NeuN in Brap^cKONPC^ cortices at 6 months. (K) Representative Gfap (green) and CD68 (red) double immunohistological images, showing increased astrogliosis and microgliosis surrounding the spongiform vacuoles (positions of vacuoles are marked by asterisks). Nuclear DNA was stained with Hoechst 33342. Bars: 100 um See also Figure S1.

We further found astrogliosis and microgliosis were markedly elevated in Brap^cKONPC^ cortical tissue in an age-dependent fashion. Increased expression of glial fibrillary acidic protein (Gfap), which represents astrocyte activation, was undetectable in the cerebral cortex of Brap^cKONPC^ mice until about 3 months of age, and it then further progressed and became remarkably high as the mutant mice aged rapidly to death (Figure 4C and D; Figure S1A). Parallel to astrogliosis was the strong increase in microgliosis in Brap^cKONPC^ cortices. As the brain’s resident innate immune cells, microglia remain in a ramified resting state in healthy brains and are activated by injury or pathological processes. Analyses with pan-microglia marker Iba and activated phagocytic microglia marker CD68 both showed that microglia in Brap^cKONPC^ cortices were de-ramified, amoeboid-like, and activated (Figure 4E). To confirm neuroinflammation, we examined the expression of several key inflammatory molecules implicated in human NDs and our RNA-seq data (Dewachter et al., 2002; Frost et al., 2019; Newcombe et al., 2018; Sarlus and Heneka, 2017), including C1q, TNFα, TGFβ1, and Trem2. All of them were upregulated significantly in the cortical tissue of 3-month old Brap^cKONPC^ mice relative to that of control mice (Figure 4F). The strong astrocyte and microglial activation along with increased inflammatory cytokines in Brap^cKONPC^ cortical tissue were in line with RNA-seq data, and they together demonstrate the neuroinflammatory and neurodegenerative brain dysfunction of Brap^cKONPC^ mice.

To ascertain Brap LOF leads to neurodegeneration, we examined the brain structure of Brap^cKONPC^ mice that survived to 6 months or older. We first performed a Golgi-Cox stain to reveal the morphology of cortical neurons. While neurons in Brap^cKONPC^ brains were grossly normal in dendritic and axonal morphology, the density of their apical dendritic spines was significantly decreased in cortical layer II/III pyramidal neurons compared to those in wild type or control mice, whereas changes in dendritic spine density were insignificant in mutant mice of younger ages (Figure 4G). These results fully agreed with the impaired synaptic transmission and neuronetwork activities in Brap^cKONPC^ mice revealed by RNA-seq, further supporting an age-dependent decline in cortical neuronal functions.

To assess whether there is an age-dependent neuronal loss in Brap^cKONPC^ mice, we examined H&E stained brain sections of a set of Brap^cKONPC^ mice (n=12) that survived to 6 months or older. We found all mutant brains showed spongiform encephalopathy in the cortical gray matter (Figure 4H). The spongiform changes were widespread, resulting in striking neuronal soma deformation and vacuolization (Figure 4I, Figure S1B). While spongiform encephalopathy is characteristic for prion diseases, it was absent in younger Brap^cKONPC^ mice or control littermates, not accompanied by the accumulation of the prion protein Prp (Figure S1C), but was associated with loss of cortical neurons (Figure 4K). These indicated that it was a non-infectious neurodegenerative lesion. Many spongiform vacuoles in Brap^cKONPC^ cortices were surrounded by reactive astrocytes and microglia (Figure 4K), suggesting their association with chronic neuroinflammation in promoting neurodegenerative pathology.

Since DSBs in Brap^cKONPC^ mice are neuronal specific, we tested whether this neuronal defect accounts for the multitude of phenotypes involving both neurons and glia in Brap^cKONPC^ mice. We generated a Brap neuronal conditional knockout line by abrogating Brap in neurons of the postnatal cortex and hippocampus with a Thy1-Cre driver (Dewachter et al., 2002)(Brap^flox Thy1Cre+;^ referred to as Brap^cKONeuron^), and asked whether Brap LOF in neurons could recapitulate the phenotype of Brap LOF in NPCs. About half of the Brap^cKONeuron^ mice (33 out of 63) were found dead by 6 months of age. Analyses of Brap^cKONeuron^ brains revealed phenotypes indistinguishable from those of the Brap^cKONPC^ mice with respect to increased DSBs, γH2A.Xub, cellular senescence, tau hyperphosphorylation, astrogliosis, and microgliosis (Figure S1D-I). This indicated that the astroglia and microglia activation in Brap^cKONPC^ mice were non-cell-autonomous responses to damaged neurons. The phenocopy of Brap^cKONeuron^ and Brap^cKONPC^ mice strongly supports the notion that sustained DSBs and senescence of cortical neurons can be a main source of neuroinflammation to induce neurodegeneration.

### Cellular senescence is associated with histone H2A ubiquitination and histone proteolysis

DSBs, if unsuccessfully repaired, can lead to several alternative consequences including oncogenesis, apoptosis, and cellular senescence. To understand the mechanism by which DSBs selectively cause cellular senescence, we studied γH2A.Xub, the unique DDR of Brap LOF (Figure 2F and K) and determined whether this histone modification is associated with the chromatin reconfiguration necessary for cells to enter and persist in the senescence state. Although γH2AX, formed by phosphorylation of the Ser-139 residue of the histone variant H2AX, is an early cellular response to DSBs, DNA damage also induces mono-ubiquitination of histone H2A, which mediates transcriptional silencing and DDR signaling (Uckelmann and Sixma, 2017; Wang et al., 2004). Thus, the co-occurrence of phosphorylation and ubiquitination of H2A or H2A variants, especially the poly-γH2A.Xub in Brap deficient cells and tissues, suggested a possibility of ubiquitin-mediated histone degradation, as global histone loss has been shown in aging and cellular senescence (Feser et al., 2010; Hu et al., 2014; Ivanov et al., 2013). In agreement, we found all major histones were progressively depleted as MEFs underwent senescence, while Brap^-/-^ MEFs lost histones more rapidly than the WT MEFs (Figure 5A). This rapid histone loss can decrease nucleosome occupancy and lead to global chromatin structure alteration that may be necessary for reinforcing and stabilizing the senescent state of Brap deficient cells.

**FIGURE 5.**
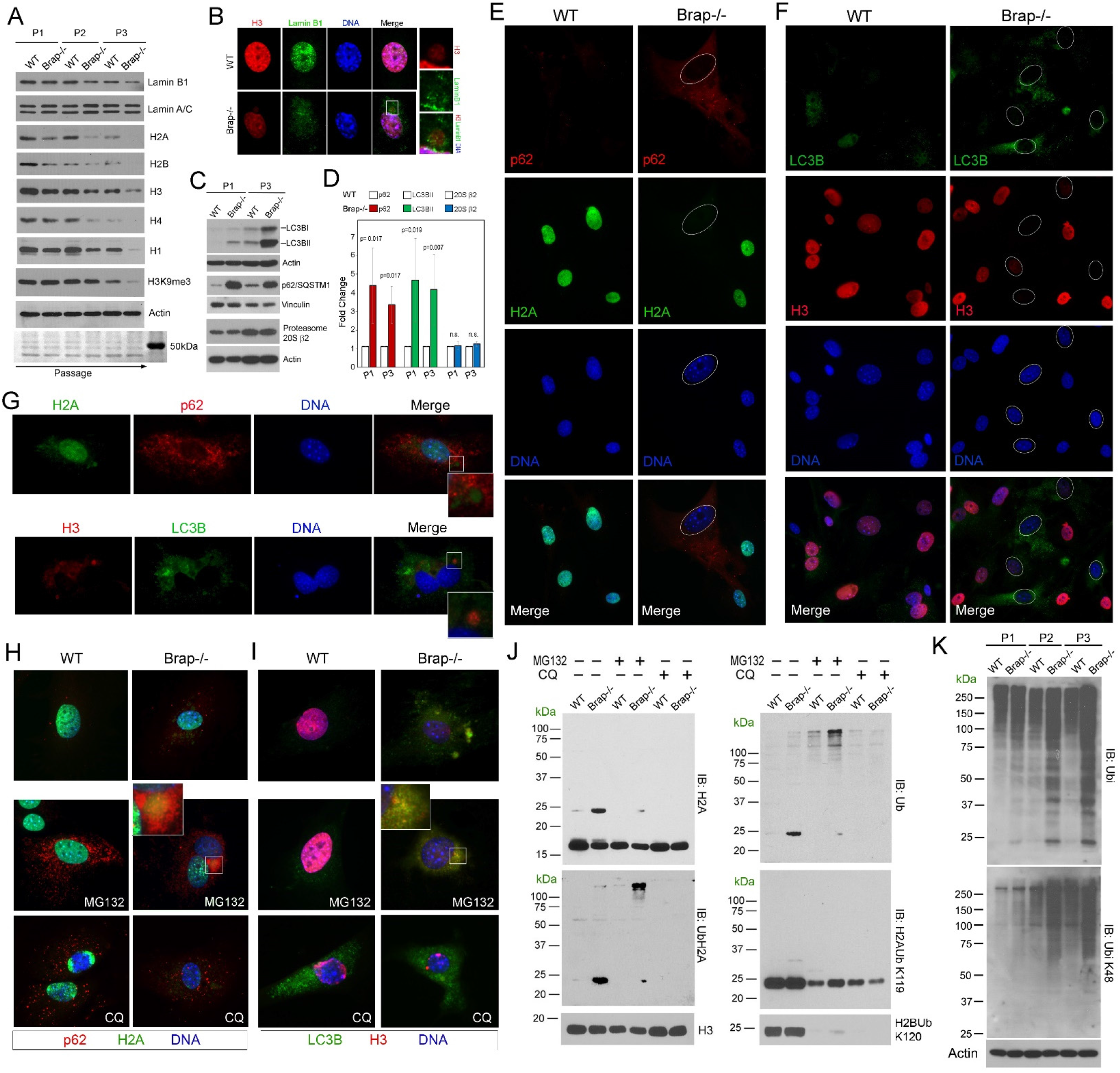
CELLULAR SENESCENCE IN BRAP^-/-^ MEFS WAS ASSOCIATED WITH HISTONE H2A UBIQUITINATION AND UPS-MEDIATED PROTEOLYSIS. (A) Immunoblotting analyses of MEF total protein extracts, showing progressive reduction of Lamin B1 and core histones with increased culture senescence. (B) Representative immunofluorescence images of histone H3 (red) and Lamin B1 (green) double stained MEFs at P3, showing the presence of cytoplasmic Lamin B1 and histone in Brap^-/-^ MEFs (boxed area at higher magnification). (C) Immunoblotting analyses of autophagy markers LC3B and p62, as well as 20S proteasome catalytic β2 subunit, showing increased autophagy flux without alteration of proteasome abundance in Brap^-/-^ relative to WT MEFs at both P1 and P3. (D) Quantification (Mean ± SD) of p62, LC3BII, and 20Sβ2 levels in WT and Brap^-/-^ MEFs at P1 and P3, respectively. n= 3-5 biological replicates. (E-G) Representative immunofluorescence images of WT and Brap^-/-^ MEFs at P3. Antibodies against p62 (red) or LC3B (green) were used to identify autophagosome and co-stained with anti-histone H2A (green) or anti-histone H3 (red). Note that in Brap^-/-^ MEFs, the reduction in nuclear histones coincides with the presence of cytoplasmic histone particles and increased p62 and LC3B but there is little cytoplasmic histone-autophagosome co-localization (boxed area at higher magnification of G). (H, I) Representative immunofluorescence images of MEFs treated with UPS blocker MG132 or lysosome blocker chloroquine (CQ), showing that cytoplasmic H2A-p62 or H3-LC3B co-localization is enhanced by MG132 but not by CQ (boxed area at higher magnification). (J) Immunoblotting analyses of histone extracts from MEFs at P2, showing Brap LOF caused elevated histone H2A mono- and poly-ubiquitination. The level of poly-H2Aub was further increased by MG132 but not by CQ, indicating UPS-mediated histone H2A proteolysis. (K) Immunoblotting analyses of MEFs total protein extracts, demonstrating increased levels of polyubiquitinated proteins in senescent Brap^-/-^ MEFs. Nuclear DNA was stained with Hoechst 33342. See also Figure S2.

The loss of nuclear histones corresponded with the detection of cytoplasmic histone particles in Brap^-/-^ MEFs. These cytoplasmic histone particles often did not contain DNA, but frequently co-occurred with the extrusion of Lamin B1 from the nucleus to the cytoplasm, while the nuclear envelope remained intact (Figure 5B, Figure S2A-C). Lamin B1’s cytoplasmic extrusion and clearance by the autophagy lysosome pathway (ALP) have been shown to mediate cellular senescence (Dou et al., 2015). Indeed, the autophagic flux was elevated in Brap^-/-^ MEFs, as evidenced by the significantly higher level of autophagy markers LC3BII and p62/SQSTM1 in Brap^-/-^ than in WT MEFs (Figure 5C and D). We also found that cells presenting stronger p62 and/or LC3B immunoreactivities were precisely those that showed reduced nuclear histones and cytoplasmic histone particles. However, immunoreactivities of cytoplasmic histone particles and p62 and/or LC3B showed little overlap (Figure 5E-G), raising a possibility that histone clearance in Brap^-/-^ MEFs was through the ubiquitin proteasome system (UPS) as suggested by the increased γH2A.Xub.

To test UPS-dependent histone proteolysis, we blocked the proteasome activity with GM132 and re-examined the cytoplasmic histone particles. We found MG132 treatment enhanced the co-localization of cytoplasmic histone with p62 and LC3B in Brap^-/-^ MEFs, while the lysosome inhibitor chloroquine (CQ) did not have this effect (Figure 5H and I). Therefore, the cytoplasmic histone particles in senescent Brap^-/-^ MEFs were destined for UPS clearance. To verify the UPS-dependent histone proteolysis, we extracted histones from Brap^-/-^ and WT MEFs and analyzed their UPS- or ALP-dependent ubiquitination state. We found that Brap LOF resulted in a specific increase in both mono- and poly-ubiquitinated histone H2A (mono- or poly-H2Aub). MG132 decreased mono-H2Aub but resulted in marked accumulation of poly-H2Aub, whereas CQ only slightly increased poly-ubiquitinated histones (Figure 5J). The coupled reduction of mono-H2Aub with elevation of poly-H2Aub upon UPS blockage suggested that mono-H2Aub in Brap^-/-^ cells is destined for poly-H2Aub and proteasomal degradation. Together, these data indicated that UPS is primarily responsible for histone H2A proteolysis in Brap^-/-^ cells, in which mono-H2Aub acts as a primer for poly-H2Aub. We did not observe changes in ubiquitin-modification of H2B, H3, H4, and H1 (Figure S2D). Although ALP-dependent histone clearance remains possible, UPS-dependent histone H2A degradation is likely the initial driver for histone degradation in Brap^-/-^ MEFs.

Histones are among the most abundant proteins in eukaryotic cells, and their increased proteolysis via UPS is likely to overwhelm the proteasome’s capacity, decreasing the clearance of other UPS substrates while stimulating the autophagic flux. Supporting an increased UPS burden in Brap^-/-^ MEFs, we found that the level and activity of proteasome 20S catalytic core complex were not significantly altered by Brap LOF (Figure 5C, D, and Figure S2E), but there was a remarkable buildup of polyubiquitinated proteins as Brap^-/-^ MEFs progressed to senescence (Figure 5K). Therefore, the depletion of histones and the accumulation of polyubiquitinated cellular proteins in Brap^-/-^ MEFs can together lead to the global epigenetic and metabolic alterations underlying the stable expression of a multitude of phenotypes that reinforce the senescence state.

### H2A ubiquitination and BRCA1 activation are the hallmark phenotypes of Brap LOF

Because H2Aub is pivotal for initiating histone proteolysis, we further evaluated whether it is a primary and early defect of Brap LOF by examining multiple tissues and ages of Brap mutant mice. We found H2Aub was significantly elevated in Brap^-/-^ and Brap^cKONPC^ mice relative to their wild type and control counterparts regardless of cell types and ages (Figure 6A; Figure S3A). Therefore, increasing the level of H2Aub is a hallmark phenotype of Brap LOF.

**FIGURE 6.**
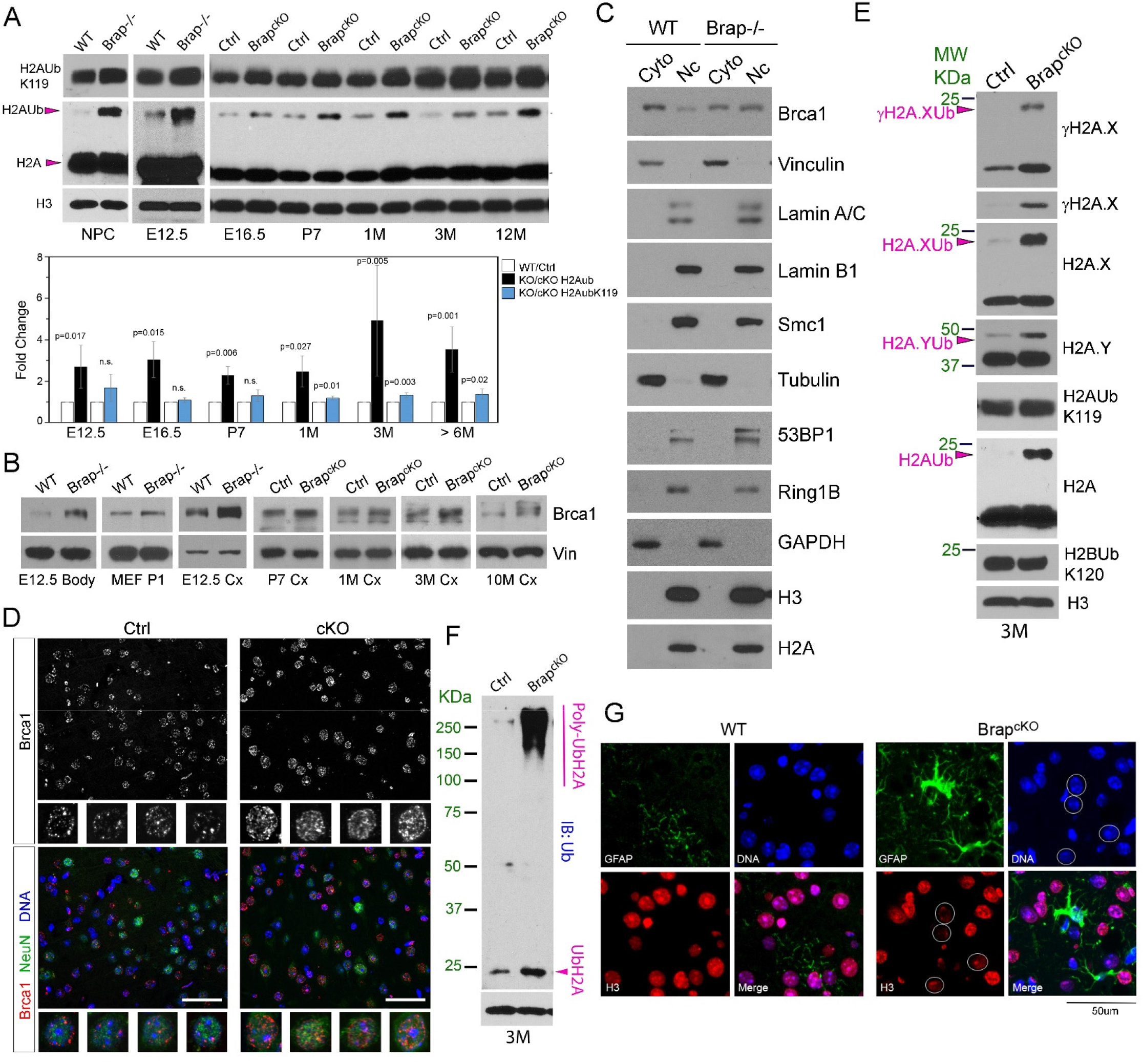
HISTONE H2A UBIQUITINATION ACCOMPANIED BY BRCA1 ACTIVATION IS THE HALLMARK PHENOTYPE OF BRAP LOF. (A) Immunoblotting of histone extracts from NPCs as well as from embryonic, neonatal, adult cerebral cortical tissues, and quantification (Mean ± SD) of increases in histone H2Aub (total H2Aub and H2AubK119, respectively) resulted from Brap LOF. n=3-6 biological replicates. (B) Immunoblotting of Brca1 in various cells and tissues, showing that Brap LOF results in increased Brca1 abundance. (C) Immunoblotting of nuclear vs. cytoplasmic fractions of MEFs at P1, showing increased nuclear localization of Brca1 in Brap^-/-^ cells. (D) Brca1 (red) and NeuN (green) double immunohistology images of cerebral cortical sections of Brap^cKONPC^ and control mice at 4 months of age. Representative images are shown. Note the increased intensity and density of Brca1 puncta in the nuclei of Brap^cKONPC^ cortical neurons (NeuN+). (E, F) Immunoblotting analyses of histone extracts from cerebral cortical tissues of 3-month old mice, showing increased ubiquitination of H2A variants targeted by Brca1 (E) along with total histone H2A ubiquitination (F). (G) Double immunohistology staining of cortical sections of 4-month old WT or Brap^cKONPC^ mice with antibodies against Gfap (green) and histone H3 (red), showing reduced nuclear histones in cells surrounded by reactive astrocytes (circles) in Brap^cKONPC^ cortical tissues. Representative images are shown. Nuclear DNA was stained with Hoechst 33342. Bars: 50 um. See also Figure S3.

Histone H2Aub can be produced by three ubiquitin E3 ligase complexes in a site-specific manner. While the ubiquitin conjugation on K119 is the most prevalent form catalyzed by the polycomb repressive complex 1 (PRC1), H2A can also be mono-ubiquitinated on K13 and K15 by RNF168/RNF8 or on K125, K127, and K129 by BRCA1 in response to DNA damage (Horn et al., 2019; Kalb et al., 2014; Tamburri et al., 2020; Uckelmann and Sixma, 2017). Our IB analyses of H2AubK119 only revealed a moderate increase in Brap deficient cells or tissues relative to controls (Figure 5J, Figure 6A, and Figure S3A). Thus, H2A in Brap deficient cells and tissues must be ubiquitinated additionally by RNF168 and/or BRCA1 on residues other than K119 due to DSBs.

BRAP was originally identified via its interaction with BRCA1, through which BRAP plays a role as a cytoplasmic docking protein to regulate BRCA1’s nuclear import. In addition, BRAP can act an E3 ligase to regulate other E3 ligases with nuclear targets (Lanctot et al., 2017; Li et al., 1998). Therefore, the phenotype of increased H2Aub due to Brap LOF is likely through elevated nuclear import and/or reduced ubiquitin-mediated turnover of Brca1. To test this, we examined the level and nuclear localization of Brca1 given their direct interaction. As expected, we found Brca1’s abundance in Brap^-/-^ MEFs, NPCs, embryos, and cortical tissues of Brap^cKONPC^ mice was consistently increased in comparison to WT and control cells or tissues (Figure 6B). In addition, we found the nuclear pool of Brca1 was much higher in Brap^-/-^ than in WT cells (Figure 6C). The high level of nuclear Brca1 was also shown by cortical neurons of the Brap^cKONPC^ mice (Figure 6D, Figure S3B). These data not only indicated that increasing Brca1’s nuclear abundance is another hallmark phenotype of Brap LOF but also suggest that the gained H2Aub in Brap deficient cells and tissues was the product of Brca1 ubiquitin E3 ligase.

The BRCA1 E3 ligase activity, though it may act independently of DSB repair, is integral to BRCA1’s role in tumor suppression (Wu et al., 2009). The substrate of BRCA1 E3 ligase include both canonical H2A and H2A variants (Kim et al., 2017). In the cortical tissue of Brap^cKONPC^ mice, we found H2A.X and H2A.Y were also ubiquitinated at higher levels relative to that of the control mice (Figure 6E, Figure S3C). This provided additional support for Brca1 as a main player for increasing H2Aub in Brap deficient cells and tissues.

Compared to the robust change in Brca1, alterations in the level and/or nuclear localization of PRC1 and RNF168 by Brap LOF were less significant. Although subtle increases in Ring1B and Bmi-1 of the PRC1 were observed in line with the moderate elevation of H2AubK119 in Brap^cKONPC^ than in control cortices, we failed to detect an upregulation of RNF168 in Brap mutants (Figure S3D and E). We also found the expression of RNF168, Ring1B, and Bmi-1 were much higher in embryonic than in mature brains, supporting their preferential requirement in brain development than in brain degeneration. Collectively, our data suggest that increased Brca1 resulting from Brap LOF is mainly responsible for H2Aub elevation.

### H2Aub is coupled with loss of proteostasis in cerebral cortical tissue and is elevated in Alzheimer’s disease

To determine whether increased histone H2Aub can drive brain aging and promote neurodegeneration, we further examined the histone content, ubiquitinated protein levels, and protein homeostasis in cortical tissues of Brap^cKONPC^ mice. Reminiscent of senescent Brap^-/-^ MEFs, increased mono-H2Aub in Brap^cKONPC^ cortices was accompanied by poly-H2Aub buildup and nuclear histone reductions in a subpopulation of cells (Figure 6F and G). We found cells showing weaker nuclear histone immuno-reactivity were surrounded by reactive astrocytes, supporting the notion that histone proteolysis stabilizes the senescent chromatin state and mediates the senescence-associated neuroinflammation. Also similar to what was observed in Brap^-/-^ MEFs, increased mono- and poly-H2Aub was accompanied by substantial elevation of the overall level of poly-ubiquitinated proteins in the Brap^cKONPC^ cortical tissue. Notably, in the cortex of Brap^cKONPC^ mice, the abundance of polyubiquitinated proteins were not obviously altered during embryonic and neonatal development, but started to elevate at young adult ages, then further built up progressively with age, and became remarkably high by 6 months of age (Figure 7A and B; Figure S4A and B). Despite the substantial backlog of poly-ubiquitinated proteins, we did not detect decreased proteasomal activities in the mutant cortical tissues. Instead, aging Brap^cKONPC^ mice showed a tendency towards elevating the catalytic activity of 20S core proteasomal complex (Figure 7C). This demonstrates clearly that the age-dependent loss of proteostasis in Brap^cKONPC^ cortices was not due to reduced UPS function but rather caused by the overproduction and backlog of polyubiquitinated proteins, among which H2Aub was the primary target resulting from Brca1-mediated DDR, while other polyubiquitinated proteins were due to UPS overflow.

**FIGURE 7.**
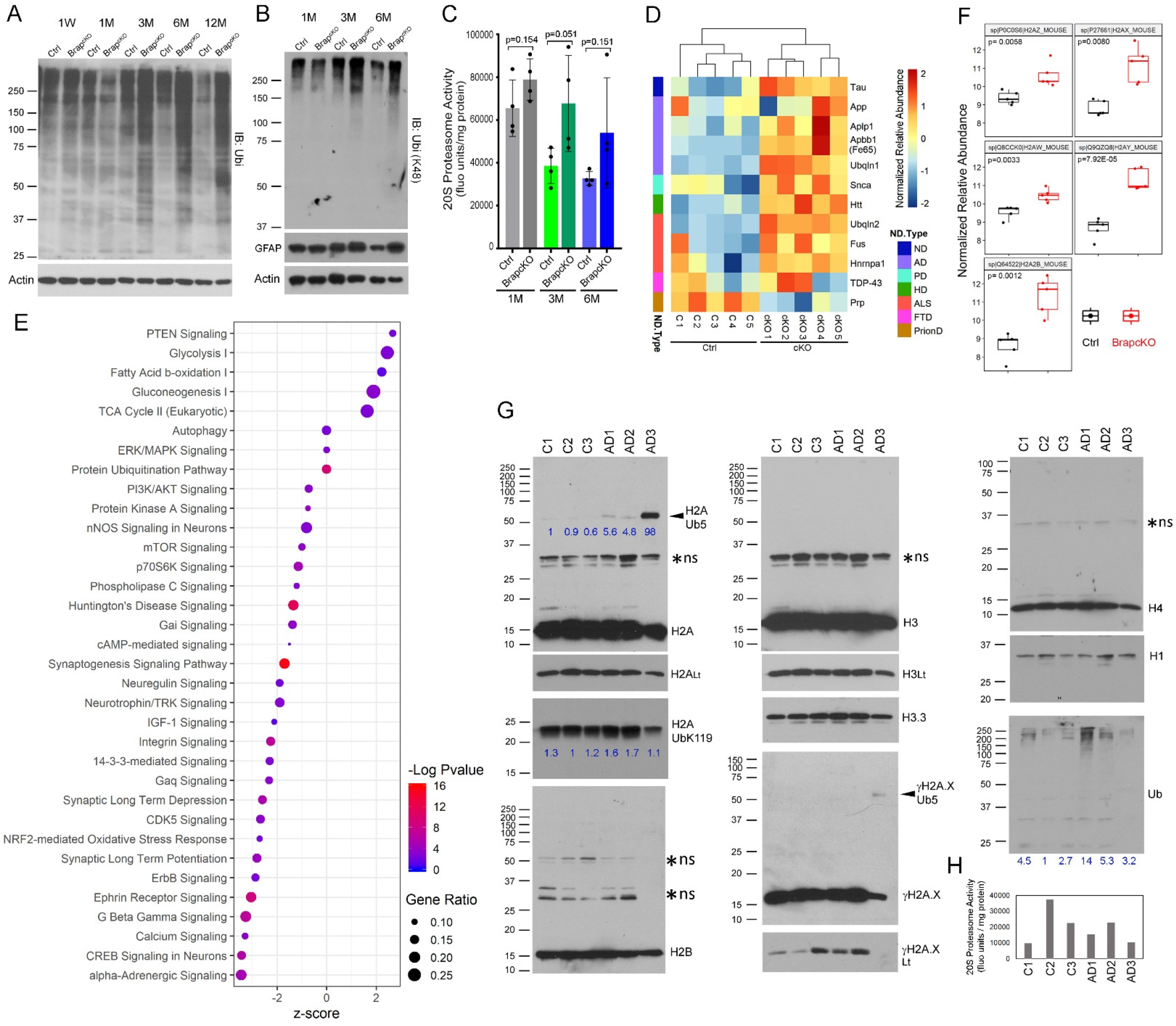
IMPAIRED PROTEOSTASIS RESULTING FROM BRAP LOF IN CEREBRAL CORTICAL TISSUE OVERLAPS WITH PROTEOPATHY OF NEURODEGENERATIVE DISORDERS. (A, B) Ubiquitin immunoblots (total ubiquitination or K48-linked polyubiquitination) of cortical tissue total protein extracts from Brap^cKONPC^ and littermate control mice, demonstrating age-dependent accumulation of poly-ubiquitinated proteins in the mutant cortical tissues. (C) 20S proteasome activities in the cortical tissue of Brap^cKONPC^ and control mice at 1 month, 3 months, and 6 months of age (Mean ± SD). (D) Heatmap of relative protein abundance shows selected proteins accumulated in aging Brap^cKONPC^ cortical tissue that are also implicated in proteopathy of human neurodegenerative disorders. ND: neurodegenerative disease; AD: Alzheimer’s disease; PD: Parkinson’s disease; HD: Huntington’s disease; ALS: Amyotrophic lateral sclerosis; PD: Prion disease. (E) Dot plot of signaling pathways altered by Brap LOF in aging cerebral cortical tissues. Shown were results of IPA of significantly altered proteins revealed by TMT analysis of Brap^cKONPC^ vs. control cortical proteome. (F) Box and whisker plots of selected results of the TMT analysis, showing significant elevation of multiple histone H2A variants in Brap^cKONPC^ relative to control cortical tissues. (G) Immunoblotting of histone extracts from postmortem cortical tissues of AD patients and age matched normal controls, showing significant elevation of H2Aub in AD cortical tissues. Asterisks denote non-specific bands (n.s.). Numbers below IB signals of H2Aub5, H2KubK119, and ubiquitinated histones denote the relative abundance normalized to histone H3. (H) 20S proteasome activities in the postmortem tissue of AD and normal control specimen being examined. See also Figure S4.

Because the progressive decline in proteostasis is a prominent hallmark of aging and a common pathological feature of neurodegeneration, we assessed the altered cortical proteome in aged Brap^cKONPC^ mice to evaluate whether it may resemble the proteopathy of NDs. We performed tandem mass tags (TMT) quantitative proteomics analysis of whole cortical proteins from 6-month old Brap^cKONPC^ and control mice. Out of the total 4708 quantified proteins, the abundance of 903 proteins was significantly altered by Brap LOF (Supplementary Table 2). Notably, many toxic proteins associated with Alzheimer’s disease (AD), Parkinson’s disease (PD), Huntington’s disease (HD), and Amyotrophic lateral sclerosis (ALS) were found to have accumulated in the Brap^cKONPC^ relative to the control cortical tissues (Figure 7D), which suggested overlapping pathogenesis between Brap LOF and human NDs. We further analyzed the alteration in proteomic profiles resulting from Brap LOF by testing their association with biological processes. In addition to aberrant protein ubiquitination and quality control, Ingenuity Pathway Analysis (IPA) showed that the cerebral cortex of Brap^cKONPC^ mice had altered glucose and fatty acid metabolism, deregulated nutrient sensing, as well as declines in numerous cell signaling pathways and neuronal network activities (Figure 7E). Notably, both long-term depression (LTD) and long-term potentiation (LTP) were significantly decreased in Brap^cKONPC^ cortices, indicating diminished synaptic plasticity and impaired cognition of the mutant brain (Figure 7E). These data demonstrated that the brain function of Brap^cKONPC^ mice was impaired at multiple levels before they die of premature brain aging and neurodegeneration.

With the remarkable proteomic alteration, multiple components of the 19S proteasome regulatory particles were significantly upregulated in Brap^cKONPC^ relative to control cortices (Figure S4C). The 19S complex recognizes ubiquitinated proteins, then deubiquitinates, unfolds, and translocates them to the 20S catalytic complex for degradation. The elevation of 19S molecules along with the tendency of increasing 20S activities in Brap^cKONPC^ mice suggested a compensatory attempt of the mutant to combat the increased UPS burden. Besides increased ubiquitination of canonical H2A, the abundance of multiple histone H2A variants, including H2A.X, H2A.Y, H2A.Z, H2A.W, and H2A.2B, was also significantly higher in Brap^cKONPC^ than in control cortices (Figure 7F). The significant changes in H2Aub and multiple histone H2A variants reflect the substantial chromatin remodeling in the Brap^cKONPC^ brain. They also support the notion that aging and neurodegeneration can be initiated by alterations of histone H2A and H2A variants.

The overlapping transcriptomic and proteomic alterations between Brap^cKONPC^ mice and NDs prompted us to determine whether the brain defect caused by Brap LOF may underlie human brain aging and neurodegeneration, as there is an urgent need for animal models to uncover the origin and recapitulate the progression of NDs. To test whether our findings were applicable to human conditions, we acquired postmortem brain samples of three Alzheimer’s disease (AD) patients along with age-matched normal controls to study their histone levels and histone ubiquitination states (Supplementary Table 3). Intriguingly, our analysis of histone extracts from the human tissue specimen demonstrated that, compared to histone H2A, H2B, H3, or H4, the level of histone H2Aub was overall higher in the cortical tissue of all three AD than in that of control individuals (Figure 7G; Figure S4D, E). The abundance of mono- and/or poly-H2Aub, albeit variable in different AD individuals and detection conditions, was notably increased in AD relative to control brains. All three AD specimens presented a marked enrichment of the unique 60 kDa Penta-ubiquitinated H2A (H2Aub5), while increases in H2AubK119 in AD brains were relatively subtle (Figure 7G). Moreover, similar to our observations of Brap^cKO^ mice, poly-ubiquitinated histones were found at higher levels in the cortical tissue of AD than in normal age-matched individuals, while a decrease in 20S proteasome activity was not obvious in AD tissues (Figure 7G, H). The close resemblance in histone ubiquitination between AD and Brap^cKO^ mice supports the relevance of our data to human NDs. It also suggests that H2Aub is a common epigenetic modification associated with brain aging and neurodegeneration, and that the Brap^cKO^ mouse model can be useful for studying AD and NDs.

## DISCUSSION

In this study, we show that Brap LOF causes accelerated aging at the cellular, tissue, and organismal levels, resulting in significant lifespan shortening in mice. This finding is consistent with the results of recent large-scale GWAS that identified the association of the *BRAP* locus with human lifespan. They together suggest that a better understanding of *BRAP*’s gain and loss of function would provide insight into the molecular determinants for aging and longevity. Moreover, our studies of the cerebral cortical NPC- and neuron-specific Brap conditional knockout mice reveal a histone H2Aub-based mechanism that connects multiple aging hallmarks to neurodegeneration, serving as a new model for aging-associated diseases.

### The interconnection of multiple aging hallmarks underpins the complex etiology of aging-associated diseases

Aging is an inevitable natural process driven by numerous molecular and cellular events that act synergistically to cause irreversible functional decline of organs involved. Through the model of Brap LOF we have learned that the abnormal increase in histone H2Aub can serve as a nexus for multiple aging accelerating mechanisms. Histone H2A mono-ubiquitination has been implicated as an evolutionarily conserved aging biomarker from an ubiquitylome study of long-lived *Drosophila* proteins. It was shown to increase in an age-dependent manner not only in *Drosophila* but also in the brain of mice, non-human primates, and humans (Yang et al., 2019). While our results partially echo this result by showing a moderate increase of H2AubK119 in Brap mutant mice, we also demonstrate that the gained H2Aub in the aging Brap^cKO^ mouse and AD brains was also outside of K119. Moreover, we show that mono-H2Aub can initiate poly-H2Aub and UPS-mediated histone degradation. The histone proteolysis not only changes chromatin structure but also increases the UPS burden, leading to proteopathy and an irreversible senescent state in cells being affected.

It is especially intriguing that the overall level of H2Aub is increased in the cortical tissue of AD. Although the terminal brain pathology of AD is characterized by the accumulation of amyloid-β and hyperphosphorylated tau proteins, the cause of idiopathic AD remains elusive, though aging is known as the major risk factor. Our finding here suggests that histone H2Aub is a potential biomarker for AD, since the level of H2Aub may mark cells’ or organs’ pathophysiological age instead of chronological age by connecting chromatin aberration to cellular senescence and proteinopathy. While H2Aub-mediated epigenetic and proteomic changes cause profound cell-intrinsic phenotypes, the senescence-associated autocrine and paracrine signaling can induce tissue-wide changes and immune-activation, leading to brain deterioration at multiple levels. Therefore, epigenetic aging H2Aub may underlie NDs and other aging-associated diseases.

### BRAP regulates BRCA1 to control H2A turnover in a context-dependent manner

Brap is a multifunctional protein cloned originally through its ability to bind to the nuclear localization signal (NLS) of the BRCA1 tumor suppressor (Li et al., 1998). As a cytoplasmic protein, BRAP can control BRCA1’s shuttling between the nucleus and the cytoplasm. BRAP is also a Ras effector and an ubiquitin E3 ligase that limits the activation of MAP kinase cascade by auto-ubiquitination (Matheny et al., 2004). In the developing central nervous system, Brap regulates MAPK signaling differentially according to the spatial position of NPCs (Lanctot et al., 2013). Brap E3 ligase can also regulate the turnover of other E3 ligases that modify many nuclear proteins (Lanctot et al., 2017). These make BRAP a versatile, yet powerful, molecule capable of connecting incoming signals that cells receive to cells’ functional outputs governed by chromatin states. Our data demonstrate BRCA1 is one of the E3 ligases regulated by BRAP. Although the role of BRCA1 in tumor suppression is attributed to the nuclear pool, variations in BRCA1’s nuclear vs. cytoplasmic localization have a strong correlation with aggressive tumor features and/or poor prognosis (Chen et al., 1995; Scully et al., 1996; Wilson et al., 1999). Nonetheless, the function of cytoplasmic BRCA1 remains unknown. We showed that the cytoplasmic pool of Brca1 was decreased upon Brap LOF. As a result, we observed higher nuclear and overall Brca1 abundance, excessive histone H2Aub, cellular senescence, and accelerated aging. This implicates an anti-senescence and anti-aging role of the cytoplasmic BRCA1. Such function of BRCA1 may be essential in stem cells and healthy somatic cells, whereas the nuclear BRCA1 pool dominates in cells with genomic lesions. This would well explain the requirement of Brca1 in development as well as the cellular senescence phenotype upon Brca1 abrogation (Cao et al., 2003; Pulvers and Huttner, 2009). Further learning the conditions by which BRAP controls BRCA1’s nuclear and cytoplasmic homeostasis is necessary to delineate the diverse roles of BRCA1 in development, aging, and oncogenesis.

Besides a major function in DSB repair, BRCA1 is well known for its ubiquitin E3 ligase function. Mutations within the N-terminal RING domain of BRCA1, which abrogates the E3 ligase activity, have been linked to breast and ovarian cancers. The BRCA1 E3 ligase activity was further found to be independent of its role in DSB repair but essential for tumor suppression (Reid et al., 2008; Wu et al., 2008; Zhu et al., 2011). Data of this study suggest that, by targeting histone H2A for ubiquitination and proteolysis, the BRCA1 E3 ligase may direct cells with irreparable DSBs to senescence. This mechanism can contribute significantly to BRCA1’s role in tumor suppression, though it may be at the expense of exhausting stem cells and speeding up the aging of those cells or organs with increased genome instability as seen in the Brap LOF model.

The BRCA1 E3 ligase modifies the C-terminus of histone H2A on K125/127/129. In Brap mutants, this activity of BRCA1 appears to promote histone degradation and nucleosome destabilization. H2A sits on the edges of the histone octamer with the flexible C-terminus tail protruding from the globular nucleosome particle at the DNA entry/exit site. Therefore, dynamic modification of H2A C-termini can have strong influences on nucleosome stability and chromatin structure. The histone H2A family contains numerous isoforms and variants, among which the highest degree of diversification is in their C-termini. It is intriguing that many histone H2A variants were increased along with H2Aub in the aging Brap^cKONPC^ brain. This could reflect the compensation for the loss of canonical H2A or alterations in the turnover of H2A variants. The end result of varying multiple H2A variants is a global chromatin structure change. Additional insights on the altered nucleosome structure as well as on the genomic loci involved in these H2A modification and variants will allow a better understanding the chromatin state that drives senescence and aging.

### The senescent states are diverse, context-dependent, and can apply to neurons

Cellular senescence, defined traditionally as a cell that can no longer divide, is essentially a stress response to reactive oxygen species, DNA damage, and protein misfolding or aggregation. However, cellular senescence presents many acquired characteristics beyond cell cycle arrests, such as chromatin remodeling, metabolic activation, and SASP. Although NPCs and MEFs with Brap deficiency undergo canonical cellular senescence in culture, the senescent phenotype in cortical tissues of Brap^cKO^ mice is more complex. Despite increases in bona fide senescence markers of SA-β-Gal and p16^Ink4a^, there was no replication arrest of any type of brain-resident cells in the Brap^cKONPC^ brain. Because the age-dependent neuroinflammation, proteopathy, brain structural deterioration, and mid-life mortality were linked to Brap LOF in neurons showing DSBs, it suggests that cellular senescence is dictated by the chromatin state. Such chromatin state not only entails the permanent cessation of propagating cells with irreparable damage but also elicits a multitude of damage responses, leading to the widespread transcriptome, metabolome, and proteome alterations for cells to adjust their microenvironment and their interaction with neighboring cells. Therefore, cellular senescence might be better defined by the distinctive chromatin state, metabolic pattern, and SASP than by cell cycle arrest alone. By this definition, neurons, which are intrinsically postmitotic, can further progress to a senescent state if they harbor sustained DSBs, leading to brain-wide metabolic and inflammatory changes.

Our RNA-seq with bulk cortical tissue identified 373 upregulated genes in Brap^cKONPC^ mice, of which 80 encode secreted proteins. While some of these transcripts could be from activated astrocytes, microglia, and lymphocytes, many are expected to represent primary SASP of Brap deficient cortical neurons with newly adopted epigenetic states. As the molecular responses of cellular senescence are context-dependent, the molecular composition of SASP is expected to be heterogeneous and pleiotropic, which can be both protective and destructive depending on the stage of phenotype progression. Further studies to identify primary SASP molecules from Brap deficient cortical neurons in young and aged mice, respectively, will provide in-depth information on the initiation and progression of neuroinflammation and degeneration.

### Intrinsic sources of genome instability in cerebral cortical neurons

DNA damage, particularly DSBs, have been shown to increase in brains of AD, PD, and ALS patients (Kim et al., 2020; Mitra et al., 2019; Schaser et al., 2019; Shanbhag et al., 2019). While genomic instability is considered a contributor in the pathogenesis of neurodegenerative disorders, the causes of neuronal DSBs and the mechanism by which DSBs induce the degenerative brain pathology remain to be defined. In the cerebral cortical tissue, Brap LOF resulted in chronic DSBs specifically in neurons. While this may be due to neurons’ high metabolic rate, potent generation of reactive oxygen species, and abundant polyunsaturated fatty acids prone to peroxidation, our data also demonstrate that neuronal DSBs could be the consequence of aberrant chromatin remodeling in neuronal differentiation and functional plasticity (Chomiak et al., 2021). During cortical neurogenesis, Brap acts synergistically with Nde1 in compacting constitutive heterochromatin composed of long tandem repetitive satellite DNA sequences flanking centromeres and telomeres. Brap LOF impairs constitutive heterochromatin remodeling and results in not only DSBs but also de-repression of satellite DNA transcription and nuclear architecture aberrations in cortical neurons (Chomiak et al., 2021). Because of the repetitive nature of satellite DNA, damages to these elements tend to perpetuate and are refractory to repair. Since neurons lack the ability of regeneration, the aberrant protection to constitutive heterochromatin is likely a main source for vulnerability to genotoxic insults throughout neurons’ lifespan. Notably, chromatin immunoprecipitation (ChIP) analyses have shown that BRCA1-mediated mono-H2Aub is enriched in the satellite regions of the genome (Padeken et al., 2019; Zhu et al., 2011). Therefore, the persistently high level of histone H2Aub seen in Brap^cKONPC^ mice may not only mark the damaged loci but also alter the structure of constitutive heterochromatin. Consequently, the broad change in chromatin state can alter gene expression and metabolism to drive neurodegeneration.

### The deep chromatin root of neurodegenerative disorders and Alzheimer’s disease

Various NDs share common defects in the accumulation and aggregation of misfolded proteins despite the fact that each of them presents unique pathological hallmarks, specific brain regional involvement, and distinctive clinical symptoms. Tremendous effort has been made to target protein aggregates, especially the amyloid plaques in AD, but the efficacy of these treatment regimens has been uncertain, suggesting that the aggregated proteins are the terminally pathological manifestations rather than the original cause of neurodegeneration. It is currently believed that AD, and many other NDs, may have a long asymptomatic or prodromal phase that lasts as long as decades, during which the diseases are frequently associated with chronic neuroinflammation and brain vascular dysfunction. The complex etiology of NDs has also made it difficult to establish proper animal models. None of the existing models of AD, PD, or ALS fully phenocopies the human disease conditions and progression. It is thus extremely challenging to identify the early diagnostic markers as well as the initial cellular and molecular defect that primes the progressive brain tissue-wide deterioration.

We believe the Brap^cKO^ mouse model will be invaluable for understanding the molecular mechanism and early-phase pathogenesis of NDs and AD. The wide range of phenotypes presented by the Brap^cKO^ mice is well in line with the disease states of human NDs. Its molecular profiles of neuroinflammation as well as gene expression and proteomic aberrations overlap substantially with hallmark pathologies of AD, PD, HD, and ALS. Therefore, data obtained from Brap^cKO^ mice are justified to serve as entry points for exploring the deep roots and complex etiology of neurodegeneration. As suggested, there is a chromatin origin of AD and NDs. Given that dynamic chromatin remodeling is an intrinsic need for neuronal plasticity, genome instability and epigenome aberrations are expected to have a constant and lifelong impact on neuronal function and longevity. A better understanding of chromatin quality control holds promises for the identification of early and effective targets and therapeutic intervention for NDs.

## STAR METHODS

### KEY RESOURCE TABLES

Supplemental Table 1: RNA-seq Data Table

Supplemental Table 2: TMT Proteomics Data Table

Supplemental Table 3: Human Postmortem AD and Control Brain Tissue Specimen Table

Antibodies Table

Supplemental Video

### EXPERIMENTAL MODEL AND SUBJECT DETAILS

#### Mice

*Brap* knockout (*Brap*^-/-^), floxed (*Brap*^flox/flox^), and Emx1-cre mediated conditional knockout (*Brap^cKONPC^*) mice have been described previously (Lanctot et al., 2017). The Thy1-cre mice were purchased from JaxMice (Stock No: 006143) and used to generate *Brap^cKONeuron^* by crossing with *Brap*^flox/flox^ mice. All mice used for this study were housed and bred according to the guidelines approved by the IACUC committees of Northwestern University and Uniformed Services University of Health Services in compliance with the AAALAC’s guidelines. Experiments were performed using littermates or age and genetic background matched control and mutant groups in both sexes. For timed matings, the day of vaginal plug was considered E0.5.

#### Cell Culture

Neural stem/progenitor cells were isolated from embryonic cortices at E12.5. Single cells were prepared and cultured in DMEM/F12 with N2 and B27 supplements, Penicillin-Streptomycin, Glutamine, Heparin, and growth factors (20 ng/ml EGF and 10 ng/mL FGF). Mouse embryonic fibroblasts (MEFs) were isolated from embryos at E12.5, after removing head and liver. The embryos were minced and digested in 0.25% trypsin-EDTA at 37°C with agitation for 10 min. Single cell suspensions were plated at a density of ≥ 10^4^/cm^2^, which is considered passage 0. The cells were then cultured according to the standard 3T3 cell protocol in DMEM supplemented with 10% FBS and Penicillin-Streptomycin. To block UPS or lysosome, 10 uM MG132, or 25mM chloroquine were applied to MEF culture for 4-12 hours.

### METHOD DETAILS

#### Immunoblotting

Immunoblotting of total cell or tissue proteins was performed by extracting with boiling 2x SDS PAGE sample buffer (62.5 mM Tris-HCl, pH 6.8, 2.5% SDS, 0.7135 M β-mercaptoethanol, 10% glycerol, 0.002% Bromophenol Blue) to fully dissolve the tissue proteins, heating at 95°C for 10 min to ensure protein denaturation, and passing through syringes with a 29^1/2^ gauge needle three times to sheer nuclear DNA and obtain homogenous extracts. 10-30 ug of total proteins were used for each immunoblotting analysis. Loadings were adjusted and normalized by the total protein content according to Coomassie blue stain of the gel after SDS PAGE and by the level of housekeeping proteins.

#### Immunostaining, Immunofluorescence, and Immunohistological Analyses

Immunofluorescence staining of cells and brain tissue sections was carried out as described (Lanctot et al., 2017; Lanctot et al., 2013). Briefly, cells were fixed with either 4% formaldehyde or cold methanol, blocked in 1% BSA and 5mg/ml lysine, and immuno-stained in a buffer containing 25 mM Hepes, pH 7.4, 250 mM Sucrose, 25 mM KCl, 25 mM Mg(CH_3_COO)_2_, 1% BSA, and 0.25% Saponin. Mouse brains were fixed by transcardial perfusion with PBS and 4% paraformaldehyde and then processed in 12 um cryosections or 5 um paraffin sections. After treating with antigen unmasking solutions (Vector Labs), brain sections were blocked with 5% goat serum and incubated with primary antibodies in PBS, 0.05% Triton X100, and 5% goat serum at 4°C overnight, and followed by staining with fluorescence conjugated antibodies and Hoechst 33342. Epifluorescence images were acquired with a Leica CTR 5500 fluorescence, DIC, and phase contrast microscope equipped with the Q Imaging Regita 2000R digital camera. Images were imported to Adobe Photoshop and adjusted for brightness and black values.

#### Senescence-associated β-gal (SA-β-gal) Staining

For identification of cellular senescence, cells were fixed with 2% formaldehyde and 0.2 % glutaraldehyde for 5 min at room temperature. Mouse brains were transcardially perfused and fixed with ice cold PBS followed by 4% paraformaldehyde and 5 mM MgCl_2_ in PBS for 6-8 hours. The fixed brains were cryopreserved and prepared as cryosections 16 um in thickness. Fixed cells or brain sections were stained with a staining solution containing 5 mM Potassium Ferrocyanide, 5 mM Potassium Ferricyanide, 150 mM NaCl, 5 mM MgCl_2_, 40 mM Citric acid-Na phosphate buffer pH 5.95, and 1 mg/ml X-gal at 37°C for 6-20 hours.

#### Golgi-Cox Staining and Dendritic Spine Analysis

Mice were euthanized with CO_2_; brains were quickly dissected, rinsed with deionized water, immersed in impregnation solution, and processed using FD Rapid GolgiStain kit (FD NeuroTechnologies) according to manufacturer’s instructions. Stained sections were examined under a Leica DM5000 light microscope. Pyramidal neurons in the cerebral cortex and hippocampus regions were imaged with a 40x objective and photographed. For dendritic spine density analysis, 16-20 pyramidal neurons in neocortical layer II/III of each mouse were randomly selected for assessment. The number of spines per 10 micrometers in secondary apical dendrites (branched from primary dendrites arising from the soma) was scored using the NIH Image J software.

#### Histone Extractions

The brain tissues used in this project were provided by the University of California Irvine Alzheimer’s Disease Research Center (UCI ADRC) and the Institute for Memory Impairments and Neurological Disorders (UCI MIND). Cell or tissues were re-suspended or homogenized and lysed in PBS containing 0.5% Triton X 100, 25 ug/ml leupeptin, 10 ug/ml Pepstatin A, 5 ug/ml Aprotinin, 10 mM Benzamidine, 2 mM PMSF, 10mM *N*-Ethylmaleimide (NEM), 10mM iodoacetamide (IAA), and 0.02% NaN_3_ at a density of ∼10^7^ cells/ml. Nuclei were first collected by centrifuge at 2500 x g for 10 min at 4°C, washed once, and re-suspended in 0.2 N HCl at a density of 4×10^7^ nuclei per ml to extract histones overnight at 4°C. After clearing debris by centrifugation at 10,000 x g for 10 min at 4°C, the histone containing supernatants were neutralized with 2 M Tris Base. Protein concentration was determined by measuring absorbance at 280 nm. Histone extractions were stored in aliquots at -20°C.

#### Cytoplasmic-nuclear Fractionation

Cells grown on culture dishes were trypsinized, collected in centrifuge tubes with DMEM and 10% FBS, and washed with PBS. After complete removal of PBS, cells were first treated gently with hypotonic buffer (10 mM Hepes, pH 7.9, 1.5 mM MgCl_2_, and 10 mM KCl) and protease inhibitors on ice for 10 min before the addition of NP-40 to a final concentration of 0.1%. After gently mixing, cells were spun at 2,500 x g for 10 min at 4°C. Supernatants were collected and analyzed as cytoplasmic fractions either directly or precipitated by 10% TCA. Pellets were gently washed twice with the hypotonic buffer by spinning at 2500 x g for 5 min before being analyzed as nuclear fractions.

#### RNA Isolation and Quantitative RT-PCR

Cerebral cortical tissue was homogenized in TRIzol reagent (Thermo Fisher) followed by total RNA extraction according to the manufacturer’s protocol. 1ug RNA was reverse transcribed into first-strand cDNA using Superscript III reverse transcriptase (Invitrogen). qRT-PCR reactions were performed using Power SYBR Green PCR Master Mix on a Roche LightCycler 480 Real-Time PCR system. Primers used for accessing gene expression were synthesized according to validated primer sequences from the MGH-PGA PrimerBank: p16Ink4a ( forward: CGCAGGTTCTTGGTCACTGT; reverse: TGTTCACGAAAGCCAGAGCG), Pai-1 (forward: 5’-TTCAGCCCTTGCTTGCCTC; reverse: ACACTTTTACTCCGAAGTCGGT), TGFβ1 (ATGTCACGGTTAGGGGCTC; reverse: GGCTTGCATACTGTGCTGTATAG), Tnf (forward: CTGGATGTCAATCAACAATGGGA; reverse: ACTAGGGTGTGAGTGTTTTCTGT), C1qc ( forward: 5’-CCCAGTTGCCAGCCTCAAT-3’; reverse: GGAGTCCATCATGCCCGTC), Trem2 (forward: 5’-CTGGAACCGTCACCATCACTC; reverse: CGAAACTCGATGACTCCTCGG), and Tbp (forward: 5’-AGAACAATCCAGACTAGCAGCA; reverse: GGGAACTTCACATCACAGCTC). Expression was normalized to TATA-binding protein (Tbp) as an internal control and results were analyzed using the 2^-(ΔΔCT)^ method.

#### RNA Sequencing Analysis of Whole Cerebral Cortical Transcriptome

Purified total RNA from whole cerebral cortical tissue of five Brap^cKONPC^ and five control (Brap^flox/floxCre-^ and Brap^flox/WTCre-^) mice at 3 months of age were processed at the University of Chicago Genomics Facility where RNA-seq library preparation was carried out. RNA libraries were sequenced using an Illumina HiSeq 4000 platform (1 x 50 bp single end and/or paired end sequencing). RNA sequencing files were transferred to the Tarbell High-performance computing cluster of Center for Research Informatics at the University of Chicago for analysis. The quality of raw sequencing data was assessed using FastQC v0.11.5. All RNA reads were first mapped to the mouse reference genome (mm10) using STAR v2.5.2b release with default parameters. Picard v2.8.1 (http://broadinstitute.github.io/picard/) was used to collect mapping metrics. The counted reads were filtered to exclude reads with identical library- and molecule barcodes. Differential gene expression analysis was performed using the DESeq2 package. Significance was defined as FDR <0.1 and absolute Fold Change ≥ 1.5.

#### 10-plex Tandem Mass Tags (TMT) Proteomic Analysis

Five Brap^cKONPC^ and five control (Brap^flox/floxCre-^ or Brap^flox/WTCre-^) mice at 6 months of age were transcardially perfused with ice cold PBS and protease inhibitors to remove high-abundant blood and serum proteins. Cerebral cortical tissue was dissected on ice, flash frozen in liquid nitrogen, and sent to the Thermo Fisher Center of Harvard Medical School where the samples were processed for 10-plex TMT analysis. MS2 spectra were searched using the SEQUEST algorithm against a Uniprot composite database derived from Mouse proteome containing its reversed complement and known contaminants. Peptide spectral matches were filtered to a 1% false discovery rate (FDR) using the target-decoy strategy combined with linear discriminant analysis. The proteins were filtered to a <1% FDR. Proteins were quantified only from peptides with a summed SN threshold of >100 and MS2 isolation specificity of 0.5.

Quantified protein were hierarchically clustered using the Euclidean distance, average linkage. Multiple Sample tests (ANOVA) with Permutation-based FDR (FDR <0.05) were performed to see significant changes among two study groups.

#### 20S Proteasome Assay

Proteasomes were extracted from mouse cortical tissue in a lysis buffer containing 50 mM HEPES, pH 7.5, 5 mM EDTA, 150 mM NaCl and 1% Triton X-100, 1 uM DTT and 2 mM ATP. Proteasome activities were determined using a 20S proteasome activity assay kit (Millipore APT280) according to manufacturer’s instructions. The amount of cleaved AMC fragment of Suc-LLVY-AMC was quantified using a CLARIOstar Plus plate reader (BMG LABTECH) at excitation (EX) = 380/emission (EM) = 460. Reaction mixtures were incubated with 10 μ lactacystin or MG132 before addition of fluorogenic substrates to ensure the specificity of the assays.

#### Quantification and Statistical Analysis

No statistical methods were used to predetermine sample size, while all experiments were performed with a minimum of three biological replicates and all cell counts were obtained from at least ten random fields. The experiments were not randomized; the investigators were not blinded to the sample allocation and data acquisition during experiments but were blinded in performing quantitative analyses of immunohistological images using the NIH Image J software.

All statistical analyses were done using GraphPad Prism 7.0 software. Data were analyzed by one-way ANOVA or unpaired two-tailed Student’s t tests for comparing differences between different genotypes. Differences were considered significant with a p value < 0.05.

## Supporting information

Antibody list

## ACKNOWLEDGMENTS

The authors wish to thank Barrington Burnett of Uniformed Services University for communication and discussion; the Genomics Facility of University of Chicago for RNA library construction and NGS analysis; the Thermo Fisher Center for Multiplexed Proteomics at the Harvard Medical School for tandem mass tag proteomic analysis; and the Biomedical Instrumentation Center of Uniformed Services University for qRT-PCR analysis. This work was supported by startup funds of Northwestern University and USUHS to Y.F. The UCI ADRC is funded by NIH/NIA Grant P30AG066519.

## AUTHOR CONTRIBUTIONS

Y.F. conceptualized the project, designed and performed the experiments, interpreted the results, and wrote the manuscript. Y.G. performed experiments; A.A.C. initiated the study and performed the experiments. Y.H. performed data analyses, C.C.L performed experiments, H.P. performed experiments. W-C.C. and J.A. performed bioinformatics data processing and analysis. X.Z. assisted with experiments. E.B. assisted with bioinformatics data analysis. E.S.M. helped with brain pathology data interpretation and the acquisition of the human AD and control brain specimens from the UCI ADRC.

## COMPETING INTERESTS

The authors declare that they have no conflict of interest.

## DISCLAIMER

The opinions, interpretations, conclusions and recommendation are those of the authors and are not necessarily endorsed by the U.S. Army, Department of Defense, the U.S. Government or the Uniformed Services University of the Health Sciences. The use of trade names does not constitute an official endorsement or approval of the use of reagents or commercial hardware or software. This document may not be cited for purposes of advertisement.

## Notes

### Competing Interest Statement

The authors have declared no competing interest.

## REFERENCES

1. Alkuraya, F.S., Cai, X., Emery, C., Mochida, G.H., Al-Dosari, M.S., Felie, J.M., Hill, R.S., Barry, B.J., Partlow, J.N., Gascon, G.G., et al. (2011). Human mutations in NDE1 cause extreme microcephaly with lissencephaly [corrected]. Am J Hum Genet 88, 536–547.

2. Aoshiba, K., Tsuji, T., Kameyama, S., Itoh, M., Semba, S., Yamaguchi, K., and Nakamura, H. (2013). Senescence-associated secretory phenotype in a mouse model of bleomycin-induced lung injury. Exp Toxicol Pathol 65, 1053–1062.

3. Bakircioglu, M., Carvalho, O.P., Khurshid, M., Cox, J.J., Tuysuz, B., Barak, T., Yilmaz, S., Caglayan, O., Dincer, A., Nicholas, A.K., et al. (2011). The essential role of centrosomal NDE1 in human cerebral cortex neurogenesis. Am J Hum Genet 88, 523–535.

4. Bushman, D.M., Kaeser, G.E., Siddoway, B., Westra, J.W., Rivera, R.R., Rehen, S.K., Yung, Y.C., and Chun, J. (2015). Genomic mosaicism with increased amyloid precursor protein (APP) gene copy number in single neurons from sporadic Alzheimer’s disease brains. Elife 4.

5. Campisi, J., and Robert, L. (2014). Cell senescence: role in aging and age-related diseases. Interdiscip Top Gerontol 39, 45–61.

6. Cao, L., Li, W., Kim, S., Brodie, S.G., and Deng, C.X. (2003). Senescence, aging, and malignant transformation mediated by p53 in mice lacking the Brca1 full-length isoform. Genes Dev 17, 201–213.

7. Capparelli, C., Chiavarina, B., Whitaker-Menezes, D., Pestell, T.G., Pestell, R.G., Hulit, J., Ando, S., Howell, A., Martinez-Outschoorn, U.E., Sotgia, F., et al. (2012). CDK inhibitors (p16/p19/p21) induce senescence and autophagy in cancer-associated fibroblasts, “fueling” tumor growth via paracrine interactions, without an increase in neo-angiogenesis. Cell Cycle 11, 3599–3610.

8. Castello, L.M., Raineri, D., Salmi, L., Clemente, N., Vaschetto, R., Quaglia, M., Garzaro, M., Gentilli, S., Navalesi, P., Cantaluppi, V., et al. (2017). Osteopontin at the Crossroads of Inflammation and Tumor Progression. Mediators Inflamm 2017, 4049098.

9. Chen, Y., Chen, C.F., Riley, D.J., Allred, D.C., Chen, P.L., Von Hoff, D., Osborne, C.K., and Lee, W.H. (1995). Aberrant subcellular localization of BRCA1 in breast cancer. Science 270, 789–791.

10. Chomiak, A.A., Lowe, C.C., Guo, Y., McDaniel, D., Pan, H., Zhou, X., Zhou, Q., Doughty, M.L., and Feng, Y. (2021). Nde1 is Required for Heterochromatin Compaction and Stability in Neocortical Neurons. bioRxiv, 2021.2006.2025.449848.

11. Choy, K.R., and Watters, D.J. (2018). Neurodegeneration in ataxia-telangiectasia: Multiple roles of ATM kinase in cellular homeostasis. Dev Dyn 247, 33–46.

12. Coppe, J.P., Desprez, P.Y., Krtolica, A., and Campisi, J. (2010). The senescence-associated secretory phenotype: the dark side of tumor suppression. Annu Rev Pathol 5, 99–118.

13. Coppe, J.P., Patil, C.K., Rodier, F., Sun, Y., Munoz, D.P., Goldstein, J., Nelson, P.S., Desprez, P.Y., and Campisi, J. (2008). Senescence-associated secretory phenotypes reveal cell-nonautonomous functions of oncogenic RAS and the p53 tumor suppressor. PLoS Biol 6, 2853–2868.

14. Coppede, F., and Migliore, L. (2010). DNA repair in premature aging disorders and neurodegeneration. Curr Aging Sci 3, 3–19.

15. Dewachter, I., Reverse, D., Caluwaerts, N., Ris, L., Kuiperi, C., Van den Haute, C., Spittaels, K., Umans, L., Serneels, L., Thiry, E., et al. (2002). Neuronal deficiency of presenilin 1 inhibits amyloid plaque formation and corrects hippocampal long-term potentiation but not a cognitive defect of amyloid precursor protein [V717I] transgenic mice. J Neurosci 22, 3445–3453.

16. Dou, Z., Xu, C., Donahue, G., Shimi, T., Pan, J.A., Zhu, J., Ivanov, A., Capell, B.C., Drake, A.M., Shah, P.P., et al. (2015). Autophagy mediates degradation of nuclear lamina. Nature 527, 105–109.

17. Feser, J., Truong, D., Das, C., Carson, J.J., Kieft, J., Harkness, T., and Tyler, J.K. (2010). Elevated histone expression promotes life span extension. Mol Cell 39, 724–735.

18. Flanagan, K.C., Alspach, E., Pazolli, E., Parajuli, S., Ren, Q., Arthur, L.L., Tapia, R., and Stewart, S.A. (2018). c-Myb and C/EBPbeta regulate OPN and other senescence-associated secretory phenotype factors. Oncotarget 9, 21–36.

19. Freund, A., Orjalo, A.V., Desprez, P.Y., and Campisi, J. (2010). Inflammatory networks during cellular senescence: causes and consequences. Trends Mol Med 16, 238–246.

20. Frost, G.R., Jonas, L.A., and Li, Y.M. (2019). Friend, Foe or Both? Immune Activity in Alzheimer’s Disease. Front Aging Neurosci 11, 337.

21. Ghosh, K., and Capell, B.C. (2016). The Senescence-Associated Secretory Phenotype: Critical Effector in Skin Cancer and Aging. J Invest Dermatol 136, 2133–2139.

22. Gorski, J.A., Talley, T., Qiu, M., Puelles, L., Rubenstein, J.L., and Jones, K.R. (2002). Cortical excitatory neurons and glia, but not GABAergic neurons, are produced in the Emx1-expressing lineage. J Neurosci 22, 6309–6314.

23. Haber, D.A. (1997). Splicing into senescence: the curious case of p16 and p19ARF. Cell 91, 555–558.

24. Hayflick, L. (1965). The Limited in Vitro Lifetime of Human Diploid Cell Strains. Exp Cell Res 37, 614–636.

25. Hernandez-Segura, A., Nehme, J., and Demaria, M. (2018). Hallmarks of Cellular Senescence. Trends Cell Biol 28, 436–453.

26. Hoang, M.L., Kinde, I., Tomasetti, C., McMahon, K.W., Rosenquist, T.A., Grollman, A.P., Kinzler, K.W., Vogelstein, B., and Papadopoulos, N. (2016). Genome-wide quantification of rare somatic mutations in normal human tissues using massively parallel sequencing. Proc Natl Acad Sci U S A 113, 9846–9851.

27. Horn, V., Uckelmann, M., Zhang, H., Eerland, J., Aarsman, I., le Paige, U.B., Davidovich, C., Sixma, T.K., and van Ingen, H. (2019). Structural basis of specific H2A K13/K15 ubiquitination by RNF168. Nat Commun 10, 1751.

28. Houlihan, S.L., and Feng, Y. (2014). The scaffold protein Nde1 safeguards the brain genome during S phase of early neural progenitor differentiation. Elife 3, e03297.

29. Hu, Z., Chen, K., Xia, Z., Chavez, M., Pal, S., Seol, J.H., Chen, C.C., Li, W., and Tyler, J.K. (2014). Nucleosome loss leads to global transcriptional up-regulation and genomic instability during yeast aging. Genes Dev 28, 396–408.

30. Ivanov, A., Pawlikowski, J., Manoharan, I., van Tuyn, J., Nelson, D.M., Rai, T.S., Shah, P.P., Hewitt, G., Korolchuk, V.I., Passos, J.F., et al. (2013). Lysosome-mediated processing of chromatin in senescence. The Journal of cell biology 202, 129–143.

31. Kakigi, R., Endo, C., Neshige, R., Kohno, H., and Kuroda, Y. (1992). Accelerated aging of the brain in Werner’s syndrome. Neurology 42, 922–924.

32. Kalb, R., Mallery, D.L., Larkin, C., Huang, J.T., and Hiom, K. (2014). BRCA1 is a histone-H2A-specific ubiquitin ligase. Cell reports 8, 999–1005.

33. Kim, B.J., Chan, D.W., Jung, S.Y., Chen, Y., Qin, J., and Wang, Y. (2017). The Histone Variant MacroH2A1 Is a BRCA1 Ubiquitin Ligase Substrate. Cell reports 19, 1758–1766.

34. Kim, B.W., Jeong, Y.E., Wong, M., and Martin, L.J. (2020). DNA damage accumulates and responses are engaged in human ALS brain and spinal motor neurons and DNA repair is activatable in iPSC-derived motor neurons with SOD1 mutations. Acta Neuropathol Commun 8, 7.

35. Kuilman, T., and Peeper, D.S. (2009). Senescence-messaging secretome: SMS-ing cellular stress. Nat Rev Cancer 9, 81–94.

36. Lanctot, A.A., Guo, Y., Le, Y., Edens, B.M., Nowakowski, R.S., and Feng, Y. (2017). Loss of Brap Results in Premature G1/S Phase Transition and Impeded Neural Progenitor Differentiation. Cell reports 20, 1148–1160.

37. Lanctot, A.A., Peng, C.Y., Pawlisz, A.S., Joksimovic, M., and Feng, Y. (2013). Spatially dependent dynamic MAPK modulation by the Nde1-Lis1-Brap complex patterns mammalian CNS. Dev Cell 25, 241–255.

38. Lasry, A., and Ben-Neriah, Y. (2015). Senescence-associated inflammatory responses: aging and cancer perspectives. Trends Immunol 36, 217–228.

39. Li, S., Ku, C.Y., Farmer, A.A., Cong, Y.S., Chen, C.F., and Lee, W.H. (1998). Identification of a novel cytoplasmic protein that specifically binds to nuclear localization signal motifs. J Biol Chem 273, 6183–6189.

40. Lodato, M.A., Rodin, R.E., Bohrson, C.L., Coulter, M.E., Barton, A.R., Kwon, M., Sherman, M.A., Vitzthum, C.M., Luquette, L.J., Yandava, C.N., et al. (2018). Aging and neurodegeneration are associated with increased mutations in single human neurons. Science 359, 555–559.

41. Lodato, M.A., Woodworth, M.B., Lee, S., Evrony, G.D., Mehta, B.K., Karger, A., Lee, S., Chittenden, T.W., D’Gama, A.M., Cai, X., et al. (2015). Somatic mutation in single human neurons tracks developmental and transcriptional history. Science 350, 94–98.

42. Lombard, D.B., Chua, K.F., Mostoslavsky, R., Franco, S., Gostissa, M., and Alt, F.W. (2005). DNA repair, genome stability, and aging. Cell 120, 497–512.

43. Lopez-Otin, C., Blasco, M.A., Partridge, L., Serrano, M., and Kroemer, G. (2013). The hallmarks of aging. Cell 153, 1194–1217.

44. Lu, T., Pan, Y., Kao, S.Y., Li, C., Kohane, I., Chan, J., and Yankner, B.A. (2004). Gene regulation and DNA damage in the ageing human brain. Nature 429, 883–891.

45. Luczak, M.W., and Zhitkovich, A. (2018). Monoubiquitinated gamma-H2AX: Abundant product and specific biomarker for non-apoptotic DNA double-strand breaks. Toxicol Appl Pharmacol 355, 238–246.

46. Madabhushi, R., Pan, L., and Tsai, L.H. (2014). DNA damage and its links to neurodegeneration. Neuron 83, 266–282.

47. Matheny, S.A., Chen, C., Kortum, R.L., Razidlo, G.L., Lewis, R.E., and White, M.A. (2004). Ras regulates assembly of mitogenic signalling complexes through the effector protein IMP. Nature 427, 256–260.

48. Mattson, M.P., and Arumugam, T.V. (2018). Hallmarks of Brain Aging: Adaptive and Pathological Modification by Metabolic States. Cell Metab 27, 1176–1199.

49. McHugh, D., and Gil, J. (2018). Senescence and aging: Causes, consequences, and therapeutic avenues. The Journal of cell biology 217, 65–77.

50. Mitra, J., Guerrero, E.N., Hegde, P.M., Liachko, N.F., Wang, H., Vasquez, V., Gao, J., Pandey, A., Taylor, J.P., Kraemer, B.C., et al. (2019). Motor neuron disease-associated loss of nuclear TDP-43 is linked to DNA double-strand break repair defects. Proc Natl Acad Sci U S A 116, 4696–4705.

51. Munoz-Espin, D., and Serrano, M. (2014). Cellular senescence: from physiology to pathology. Nat Rev Mol Cell Biol 15, 482–496.

52. Musi, N., Valentine, J.M., Sickora, K.R., Baeuerle, E., Thompson, C.S., Shen, Q., and Orr, M.E. (2018). Tau protein aggregation is associated with cellular senescence in the brain. Aging Cell 17, e12840.

53. Newcombe, E.A., Camats-Perna, J., Silva, M.L., Valmas, N., Huat, T.J., and Medeiros, R. (2018). Inflammation: the link between comorbidities, genetics, and Alzheimer’s disease. J Neuroinflammation 15, 276.

54. Padeken, J., Zeller, P., Towbin, B., Katic, I., Kalck, V., Methot, S.P., and Gasser, S.M. (2019). Synergistic lethality between BRCA1 and H3K9me2 loss reflects satellite derepression. Genes Dev 33, 436–451.

55. Panier, S., and Boulton, S.J. (2014). Double-strand break repair: 53BP1 comes into focus. Nat Rev Mol Cell Biol 15, 7–18.

56. Pazolli, E., Alspach, E., Milczarek, A., Prior, J., Piwnica-Worms, D., and Stewart, S.A. (2012). Chromatin remodeling underlies the senescence-associated secretory phenotype of tumor stromal fibroblasts that supports cancer progression. Cancer Res 72, 2251–2261.

57. Pulvers, J.N., and Huttner, W.B. (2009). Brca1 is required for embryonic development of the mouse cerebral cortex to normal size by preventing apoptosis of early neural progenitors. Development 136, 1859–1868.

58. Quelle, D.E., Zindy, F., Ashmun, R.A., and Sherr, C.J. (1995). Alternative reading frames of the INK4a tumor suppressor gene encode two unrelated proteins capable of inducing cell cycle arrest. Cell 83, 993–1000.

59. Rao, S.G., and Jackson, J.G. (2016). SASP: Tumor Suppressor or Promoter? Yes! Trends Cancer 2, 676–687.

60. Reid, L.J., Shakya, R., Modi, A.P., Lokshin, M., Cheng, J.T., Jasin, M., Baer, R., and Ludwig, T. (2008). E3 ligase activity of BRCA1 is not essential for mammalian cell viability or homology-directed repair of double-strand DNA breaks. Proc Natl Acad Sci U S A 105, 20876–20881.

61. Rodier, F., and Campisi, J. (2011). Four faces of cellular senescence. The Journal of cell biology 192, 547–556.

62. Rulten, S.L., and Caldecott, K.W. (2013). DNA strand break repair and neurodegeneration. DNA Repair (Amst) 12, 558–567.

63. Rutten, B.P., Schmitz, C., Gerlach, O.H., Oyen, H.M., de Mesquita, E.B., Steinbusch, H.W., and Korr, H. (2007). The aging brain: accumulation of DNA damage or neuron loss? Neurobiol Aging 28, 91–98.

64. Salminen, A., Kauppinen, A., and Kaarniranta, K. (2012). Emerging role of NF-kappaB signaling in the induction of senescence-associated secretory phenotype (SASP). Cell Signal 24, 835–845.

65. Sarlus, H., and Heneka, M.T. (2017). Microglia in Alzheimer’s disease. The Journal of clinical investigation 127, 3240–3249.

66. Schaser, A.J., Osterberg, V.R., Dent, S.E., Stackhouse, T.L., Wakeham, C.M., Boutros, S.W., Weston, L.J., Owen, N., Weissman, T.A., Luna, E., et al. (2019). Alpha-synuclein is a DNA binding protein that modulates DNA repair with implications for Lewy body disorders. Scientific reports 9, 10919.

67. Scully, R., Ganesan, S., Brown, M., De Caprio, J.A., Cannistra, S.A., Feunteun, J., Schnitt, S., and Livingston, D.M. (1996). Location of BRCA1 in human breast and ovarian cancer cells. Science 272, 123–126.

68. Shanbhag, N.M., Evans, M.D., Mao, W., Nana, A.L., Seeley, W.W., Adame, A., Rissman, R.A., Masliah, E., and Mucke, L. (2019). Early neuronal accumulation of DNA double strand breaks in Alzheimer’s disease. Acta Neuropathol Commun 7, 77.

69. Soares, J.P., Cortinhas, A., Bento, T., Leitao, J.C., Collins, A.R., Gaivao, I., and Mota, M.P. (2014). Aging and DNA damage in humans: a meta-analysis study. Aging (Albany NY) 6, 432–439.

70. Suberbielle, E., Sanchez, P.E., Kravitz, A.V., Wang, X., Ho, K., Eilertson, K., Devidze, N., Kreitzer, A.C., and Mucke, L. (2013). Physiologic brain activity causes DNA double-strand breaks in neurons, with exacerbation by amyloid-beta. Nat Neurosci 16, 613–621.

71. Tamburri, S., Lavarone, E., Fernandez-Perez, D., Conway, E., Zanotti, M., Manganaro, D., and Pasini, D. (2020). Histone H2AK119 Mono-Ubiquitination Is Essential for Polycomb-Mediated Transcriptional Repression. Mol Cell 77, 840–856 e845.

72. Thadathil, N., Hori, R., Xiao, J., and Khan, M.M. (2019). DNA double-strand breaks: a potential therapeutic target for neurodegenerative diseases. Chromosome Res 27, 345–364.

73. Thiriet, C., and Hayes, J.J. (2005). Chromatin in need of a fix: phosphorylation of H2AX connects chromatin to DNA repair. Mol Cell 18, 617–622.

74. Timmers, P.R., Mounier, N., Lall, K., Fischer, K., Ning, Z., Feng, X., Bretherick, A.D., Clark, D.W., e, Q.C., Agbessi, M., et al. (2019). Genomics of 1 million parent lifespans implicates novel pathways and common diseases and distinguishes survival chances. Elife 8.

75. Todaro, G.J., and Green, H. (1963). Quantitative studies of the growth of mouse embryo cells in culture and their development into established lines. The Journal of cell biology 17, 299–313.

76. Uckelmann, M., and Sixma, T.K. (2017). Histone ubiquitination in the DNA damage response. DNA Repair (Amst) 56, 92–101.

77. Vaughan, D.E., Rai, R., Khan, S.S., Eren, M., and Ghosh, A.K. (2017). Plasminogen Activator Inhibitor-1 Is a Marker and a Mediator of Senescence. Arterioscler Thromb Vasc Biol 37, 1446–1452.

78. Wang, H., Wang, L., Erdjument-Bromage, H., Vidal, M., Tempst, P., Jones, R.S., and Zhang, Y. (2004). Role of histone H2A ubiquitination in Polycomb silencing. Nature 431, 873–878.

79. Weidenheim, K.M., Dickson, D.W., and Rapin, I. (2009). Neuropathology of Cockayne syndrome: Evidence for impaired development, premature aging, and neurodegeneration. Mech Ageing Dev 130, 619–636.

80. White, R.R., and Vijg, J. (2016). Do DNA Double-Strand Breaks Drive Aging? Mol Cell 63, 729–738.

81. Wilson, C.A., Ramos, L., Villasenor, M.R., Anders, K.H., Press, M.F., Clarke, K., Karlan, B., Chen, J.J., Scully, R., Livingston, D., et al. (1999). Localization of human BRCA1 and its loss in high-grade, non-inherited breast carcinomas. Nat Genet 21, 236–240.

82. Wu, J., Huen, M.S., Lu, L.Y., Ye, L., Dou, Y., Ljungman, M., Chen, J., and Yu, X. (2009). Histone ubiquitination associates with BRCA1-dependent DNA damage response. Mol Cell Biol 29, 849–860.

83. Wu, W., Koike, A., Takeshita, T., and Ohta, T. (2008). The ubiquitin E3 ligase activity of BRCA1 and its biological functions. Cell Div 3, 1.

84. Yang, L., Ma, Z., Wang, H., Niu, K., Cao, Y., Sun, L., Geng, Y., Yang, B., Gao, F., Chen, Z., et al. (2019). Ubiquitylome study identifies increased histone 2A ubiquitylation as an evolutionarily conserved aging biomarker. Nat Commun 10, 2191.

85. Zhu, Q., Pao, G.M., Huynh, A.M., Suh, H., Tonnu, N., Nederlof, P.M., Gage, F.H., and Verma, I.M. (2011). BRCA1 tumour suppression occurs via heterochromatin-mediated silencing. Nature 477, 179–184.

