## Supplementary Figures for "Histone H2A ubiquitination resulting from Brap loss of function connects multiple aging hallmarks and accelerates neurodegeneration"

<u>Supplemental Figure S1</u>

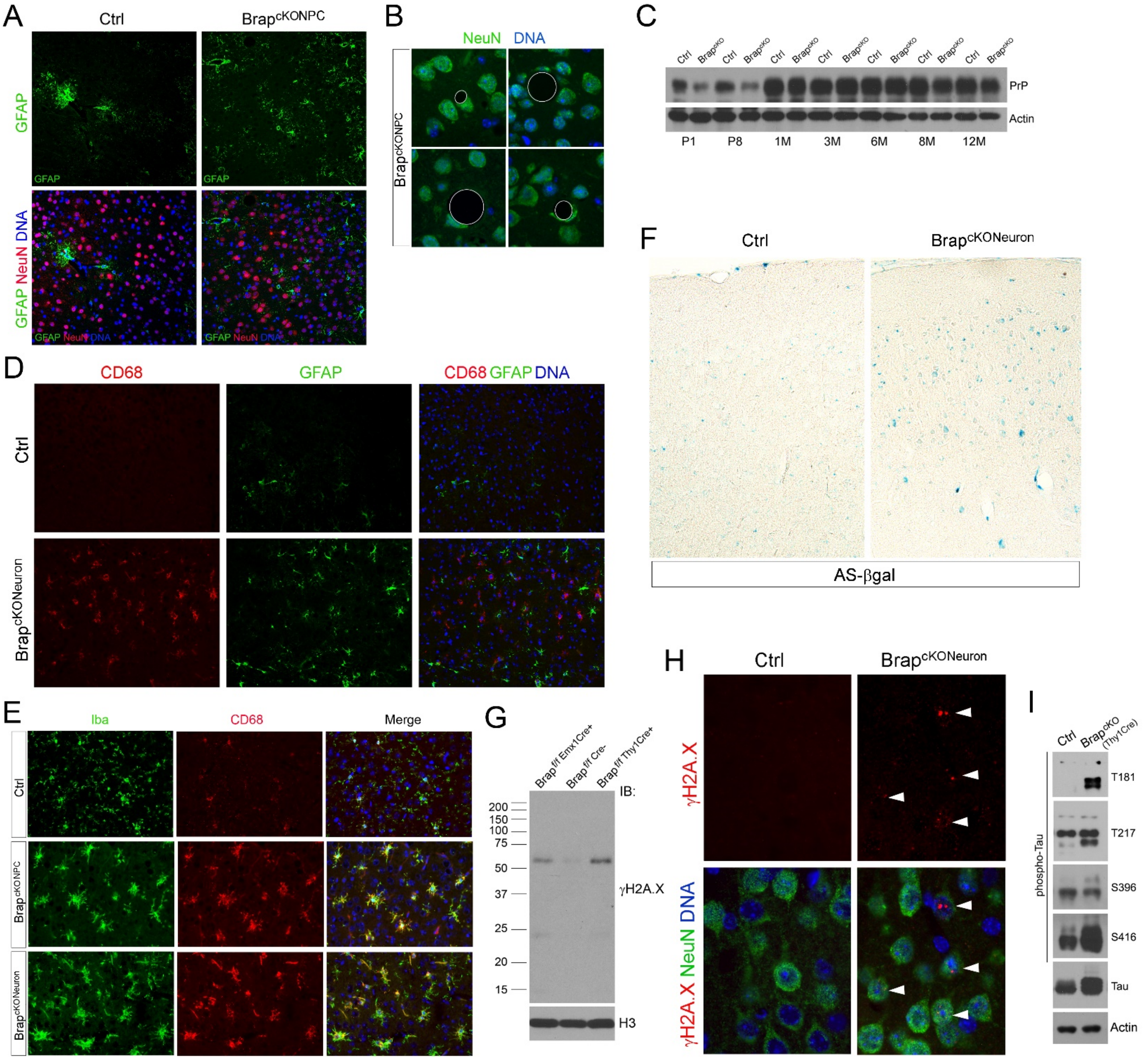

**FIGURE S1 (RELATED TO FIGURE 4). PHENOCOPY OF BRAP$^{CKONEURON}$ WITH BRAP$^{CKONPC}$ MICE**

(A) NeuN (red) and Gfap (green) double immunohistology (IH) staining of cerebral cortical sections of Brap$^{cKONPC}$ and control mice at 6 months of age. Representative images are shown.

(B) Representative images of anti-NeuN (green) immunohistological staining of cerebral cortical sections from Brap$^{cKONPC}$ mice at 10 months of age, showing deformed neurons adjacent to spongiform vacuoles (marked by circles).

(C) Immunoblotting cortical total protein extract with prion protein (Prp) antibodies of Brap$^{cKONPC}$ and control mice at various ages.

(D) Representative CD68 (red) and Gfap (green) double immunohistological images, showing microgliosis and astrogliosis in the cortical tissue of Brap$^{cKONeuron}$ mice at 3 months of age.

(E) Representative Iba (green) and CD68 (red) double immunohistological images, showing the phenocopy of Brap$^{cKONeuron}$ and Brap$^{cKONPC}$ mice with respect to microglia activation in the cerebral cortical tissue at 3 months of age.

(F) Senescence-associated β−gal analysis of cerebral cortices of Brap$^{cKONeuron}$ and control mice at 3 months of age. Representative images are shown.

(G) Immunoblotting of cerebral cortical histone extracts, showing the phenocopy of Brap$^{cKONeuron}$ and Brap$^{cKONPC}$ mice with respect to the elevation of ubiquityl- γH2A.X.

(H) NeuN (green) and γH2A.X (red) double immunohistological images of cortical section of Brap$^{cKONeuron}$ and control mice at 3 months of age, showing the presence of neuronal-specific DSBs in Brap$^{cKONeuron}$ mice.

(I) Immunoblotting of cerebral cortical total protein extracts of 3-month old Brap$^{cKONeuron}$ and control mice with anti-phospho-tau antibodies, showing tau hyper-phosphorylation on multiple residues in mutant cortices.

Nuclear DNA was stained with Hoechst 33342.

Supplemental Figure S2

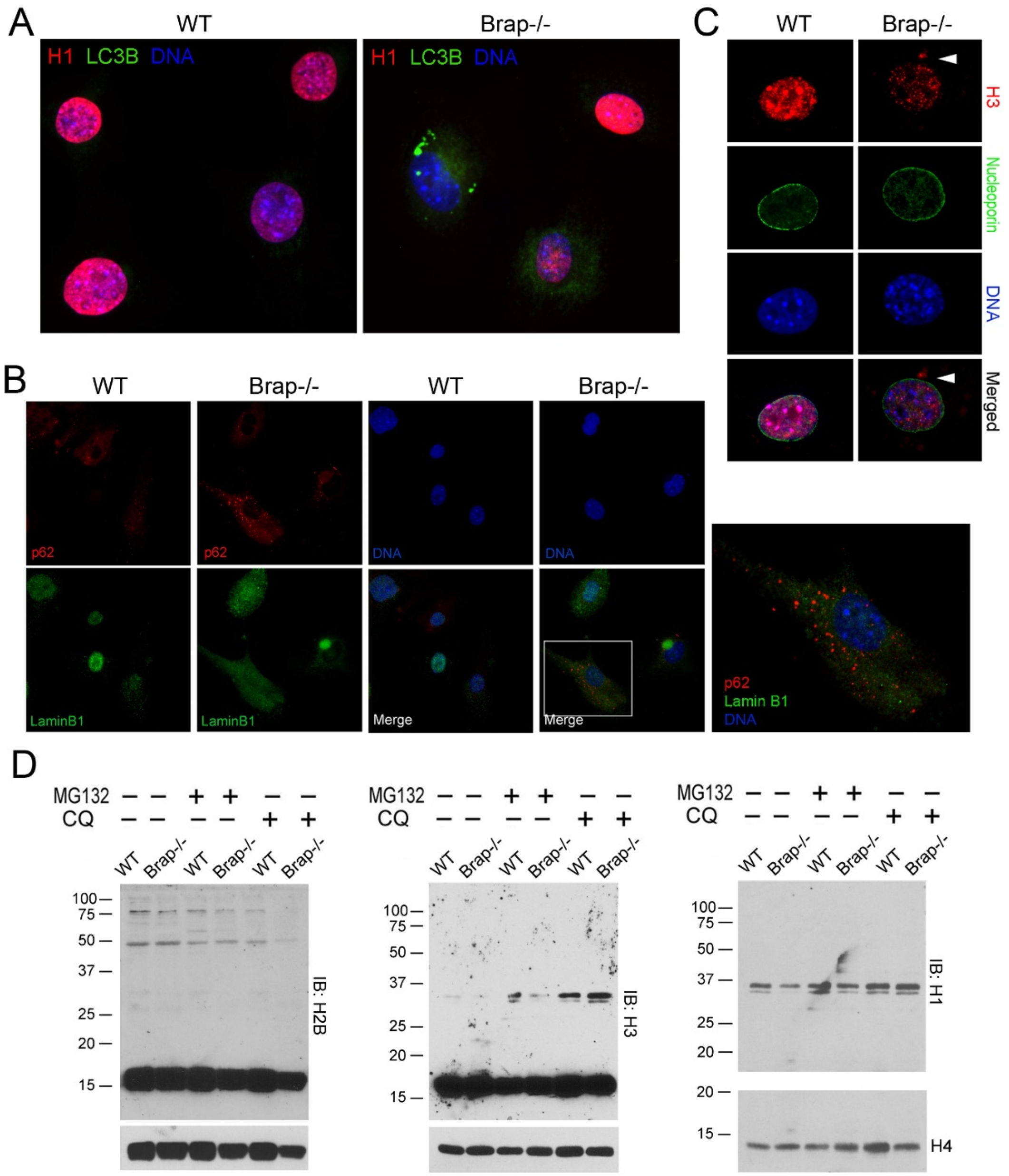


**FIGURE S2 (RELATED TO FIGURE 5). BRAP LOF RESULTS IN LOSS OF NUCLEAR HISTONES AND NUCLEAR LAMIN B1.**

(A) Immunofluorescence images of WT and Brap$^{-/-}$ MEFs at P3 stained with antibodies to histone H1 and anti-LC3B, showing the correlation of reduced histone H1 with increased cytoplasmic LC3B.

(B) Immunofluorescence images of WT and Brap$^{-/-}$ MEFs at P3 stained with antibodies against Lamin B1 and p62, showing the correlation of reduced nuclear Lamin B1 with increased cytoplasmic Lamin B1 and p62.

(C) Immunofluorescence images of WT and Brap$^{-/-}$ MEFs at P3 stained with anti-nuceloporin and histone H3, showing intact nuclear envelope despite the presence of cytoplasmic histone H3 in Brap$^{-/-}$ MEFs.

(D) Immunoblotting analyses of histone extracts from MEFs at P2, showing that the ubiquitination of histone H2B, H3, H4, and H1 was not obviously altered by Brap LOF. E. 20S proteasome activity of MEFs at P3, (Mean ± SD).

Supplemental Figure S3

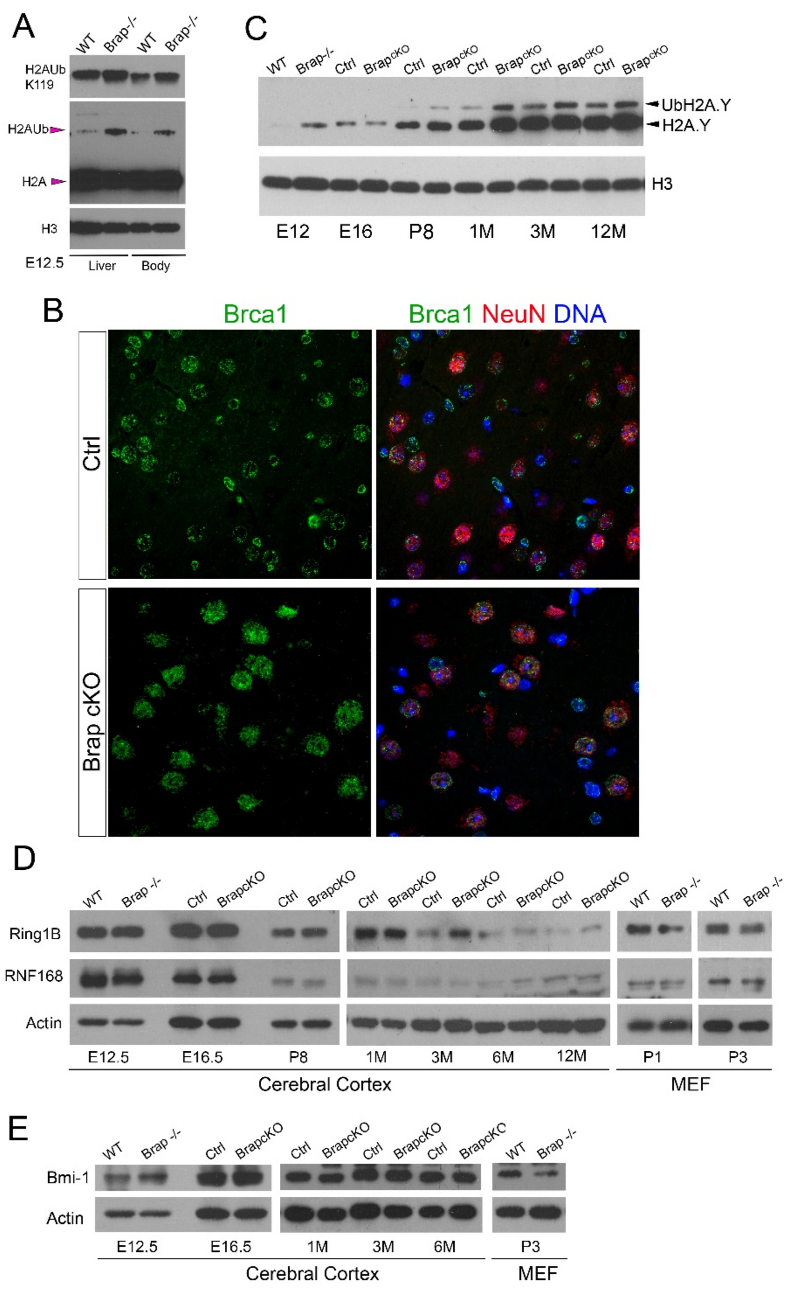


**FIGURE S3 (RELATED TO FIGURE 6). BRAP LOF INCREASES HISTONE H2AUB ALONG WITH UBIQUITYL-H2A.Y TARGETED BY BRCA1 E3 LIGASE.**

(A) Immunoblotting of histone extracts from embryonic liver and body tissues, showing higher level of H2Aub in Brap$^{-/-}$ than in of WT mice.

(B) Immunoblotting of histone extracts from cerebral cortical tissue at various ages, showing increased H2A.Y (MacroH2A) ubiquitination in postnatal, adult, and aged Brap$^{cKONPC}$ mice.

(C) Brca1 (green) and NeuN (red) double immunohistology images of cerebral cortical sections of Brap$^{cKONPC}$ and control mice at 1 month of age. Representative images are shown.

(D, E) Immunoblotting of total protein extracts from cortical tissues or MEFs from WT, Brap$^{-/-}$, Brap$^{cKONPC}$, and control mice at various ages.

Supplemental Figure S4

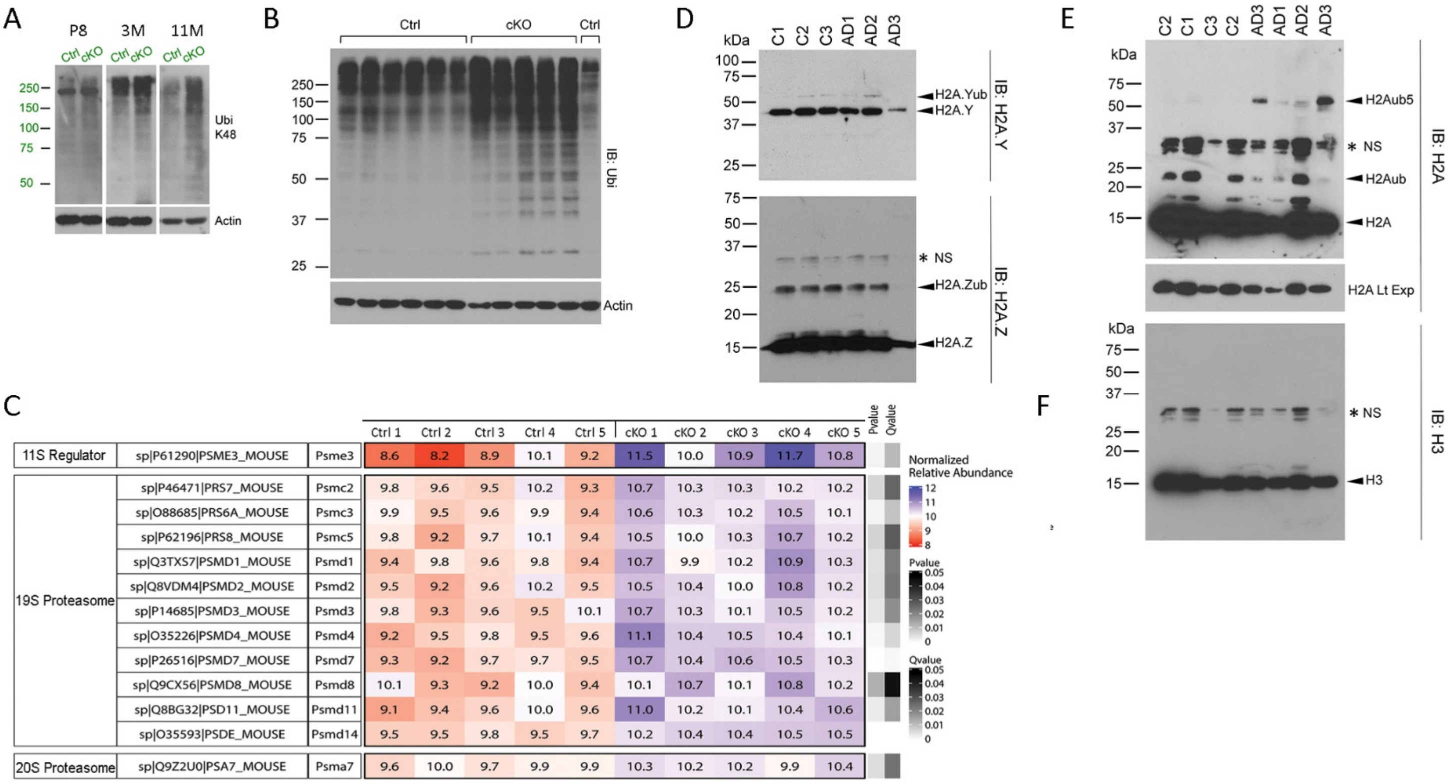


**FIGURE S4 (RELATED TO FIGURE 7). ACCELERATED BRAIN AGING OF BRAPC$^{KONPC}$ MICE IS ASSOCIATED WITH AN ACCUMULATION OF POLY-UBIQUITINATED PROTEINS IN CEREBRAL CORTICAL TISSUES ALONG WITH A COMMON INCREASE OF H2AUB IN BOTH BRAPC$^{KONPC}$ MICE AND AD PATIENTS.**

(A) Immunoblotting of cerebral cortical total protein extracts, showing an age-dependent increase in K48-linked polyubiquitinated proteins that are destined for UPS degradation in Brap$^{cKONPC}$ cortices.

(B) Anti-ubiquitin immunoblotting of total cerebral cortical protein extracts from Brap$^{cKONPC}$ and control mice at 6 months of age, showing a backlog of polyubiquitinated proteins in all Brap$^{cKONPC}$ mice.

(C) Table and heatmap, showing the significant accumulation of proteasome catalytic and regulatory proteins in Brap$^{cKONPC}$ cortical tissue revealed by TMT analysis.

(D) Immunoblotting AD postmortem cortical tissue histone extracts with antibodies against histone H2A.Y and H2A.Z, respectively. Asterisks denote non-specific bands.

(E) Immunoblotting AD postmortem cortical tissue histone extracts with antibodies against histone H2A. This represents a technical replication of Figure 7F. Asterisks denote non-specific bands.
